# Phosphorylation of the CAMTA3 transcription factor triggers its destabilization and nuclear export

**DOI:** 10.1101/825323

**Authors:** Xiyuan Jiang, Wolfgang Hoehenwarter, Dierk Scheel, Justin Lee

## Abstract

The calmodulin-binding transcription activator 3 (CAMTA3) is a repressor of immunity-related genes but an activator of cold-induced genes in plants. Post-transcriptional or -translational mechanisms have been proposed to control CAMTA3’s role in the crosstalk between immune and chilling responses. Here, we show that treatment with the bacterial flg22 elicitor, but not cold stress, induces a phospho-mobility shift of CAMTA3 proteins. Correspondingly, CAMTA3 is directly phosphorylated by two flg22-responsive mitogen-activated protein kinases (MAPKs), MPK3 and MPK6, which triggers CAMTA3 nuclear export and destabilization. SR1IP1, a substrate E3 ubiquitin ligase adaptor required for pathogen-induced CAMTA3 degradation, is shown here to be likely plasma-membrane-localized and therefore cannot physically interact with the nuclear CAMTA3. Despite the flg22-inducible re-localization of CAMTA3 to the cytoplasm, we failed to detect CAMTA3-SR1IP1 complexes. Hence, the role of SR1IP1 for CAMTA3 degradation needs to be re-evaluated. Surprisingly, flg22 elicitation can still induce nuclear export and phospho-mobility shift of a phospho-null CAMTA3 that cannot be phosphorylated by MAPKs, suggesting the participation of additional flg22-responsive kinase(s). A constitutively-active calcium-dependent protein kinase, CPK5, can stimulate a phospho-mobility shift in CAMTA3 similar to that induced by flg22. Although CPK5 can interact with CAMTA3, it did not directly phosphorylate CAMTA3, suggesting the requirement of a still unidentified downstream kinase or additional components. Overall, at least two flg22-responsive kinase pathways target CAMTA3 to induce degradation that presumably serves to remove CAMTA3 from target promoters and de-repress expression of defence genes.

**One sentence summary:** Treatment with flg22 activates two independent kinase pathways that effect CAMTA3 phosphorylation and degradation.

## Introduction

Unlike animals, plants do not possess a circulatory adaptive immune system. Instead, they rely on an efficient cellular innate immunity system to defend against various pathogens. The first layer of this immunity system is initiated by perception of pathogen- or microbe-associated molecular patterns (PAMPs/MAMPs) by pattern recognition receptors. A well-studied system is the recognition of the bacterial flagellin-derived flg22 peptide PAMP by the FLS2 receptor (Gómez-Gómez and Boller, 2000; Chinchilla et al., 2007). Ligand recognition and binding trigger receptor activation (Zipfel et al., 2004; Chinchilla et al., 2007; Roux et al., 2011) and transduce this into intracellular signaling events, including a rise of cytosolic Ca^2+^ level (Ranf et al., 2011), production of extracellular reactive oxygen species (ROS) (Kadota et al., 2014; Li et al., 2014) and activation of mitogen-activated protein kinase (MAPK) cascades (Asai et al., 2002) and calcium-dependent protein kinases (CDPKs) (Boudsocq et al., 2010). This signaling network induces transcriptional and metabolic reprogramming to establish pattern-triggered immunity (PTI). A second layer of immunity is the so-called effector-triggered immunity (ETI) that is activated after direct or indirect recognition of pathogen effectors or their modifications of host proteins, respectively. However, a distinction between PTI and ETI is not always clear (Thomma et al., 2011).

In *Arabidopsis thaliana*, the best studied stress-responsive MAPKs are MPK3, MPK4, and MPK6, which are known to be activated during pathogen infection or elicitation with PAMPs, such as flg22 and elf18 (Asai et al., 2002). An MPK4 paralog, MPK11, was subsequently identified as a fourth PAMP-activated MAPK (Eschen-Lippold et al., 2012). The activated MAPKs phosphorylate a variety of substrates to transduce upstream signals to trigger further defense responses. These substrates could be transcriptional/translational regulators, structural components or enzymes. Phosphorylation may affect the function of the substrates through altered protein stability, cellular localization, enzyme activity, or the ability to bind to other partners (Lee et al., 2015; Zhang et al., 2016). Therefore, identifying the MAPK targets responsible for regulating the downstream immunity activation is essential for engineering pathogen resistance (Hoehenwarter et al., 2012; Lassowskat et al., 2014; Rayapuram et al., 2014).

Ca^2+^ influx is one of the earliest signaling events after PAMP perception and plays a crucial role in PTI (Ranf et al., 2011). The stimulus-specific spatio-temporal features of Ca^2+^ signals are referred to as Ca^2+^ signatures (Cheval et al., 2013). Ca^2+^ sensing proteins, such as calmodulin (CaM), CaM-like proteins (CMLs), Ca^2+^-dependent protein kinases (CDPKs) and calcineurin B-like proteins (CBLs) (DeFalco et al., 2010), read and transduce these Ca^2+^ signatures into downstream cellular responses (Seybold et al., 2014). While CBLs need to recruit additional partners for further signal transduction, CDPKs are both sensor and signal relay in one entity. CDPKs are calcium-regulated serine/threonine kinases and unique to plants (Boudsocq and Sheen, 2013). Four Arabidopsis CDPKs, i.e. CPK4, CPK5, CPK6 and CPK11, are transiently activated upon flg22 perception and act as positive regulators in PTI (Boudsocq et al., 2010). CaM is highly conserved in eukaryotes, and seven CaM isoforms are encoded in the Arabidopsis genome (Ranty et al., 2006). A conformational change of CaM induced by Ca^2+^ binding can promote interaction with downstream targets that contain CaM-binding domain (CaMBD), such as transcription factors, kinases or metabolic enzymes (DeFalco et al., 2010; Cheval et al., 2013).

The Arabidopsis calmodulin-binding transcription activator 3 (CAMTA3, AT2G22300, also abbreviated as SR1) belongs to a six-membered gene family (Finkler et al., 2007). Key conserved domains include an N-terminal CG-1 domain that binds a conserved 6-bp motif (with the consensus sequence (A/C/G)CGCG(G/T/C)) in the promoter of target genes (Yang and Poovaiah, 2002) and two IQ motifs and a CaMBD near the C-terminus, which are thought to mediate Ca^2+^-independent and - dependent calmodulin binding, respectively (Finkler et al., 2007; Du et al., 2009). Transcriptomics analysis of two T-DNA insertion knockout mutants (*camta3-1* and *camta3-2*) revealed up-regulation of many defense genes (e.g. *PRs*, *NDR1*, *PAD4*, *ZAT10* and various *WRKYs*) (Galon et al., 2008). Under normal growth conditions (at 19-21°C), both *camta3* mutants display reduced growth, chlorotic lesions, constitutive expression of *PR* genes (systemic acquired resistance(SAR)-associated marker genes) and enhanced resistance to bacterial and fungal pathogens. Hypersensitive response and SAR-related features such as accumulation of SA, ROS and autofluorescent compounds are also enhanced (Galon et al., 2008; Du et al., 2009; Nie et al., 2012). Thus, CAMTA3 is thought to be a suppressor of defense responses in Arabidopsis, by directly binding to the promoter of target genes at specific “CGCG”-containing *cis*-elements. In agreement, chromatin-immunoprecipitation assay showed CAMTA3 binding to the promoters of defense regulators such as *EDS1* (*Enhanced Disease Susceptibility 1*, a positive regulator of SA biosynthesis) (Du et al., 2009), *NDR1* (*non-race-specific disease resistance 1*) and *EIN3* (*ethylene insensitive 3*) (Nie et al., 2012). During pathogen infection, repressor functions of CAMTA3 is alleviated through its degradation. This proteasome-dependent degradation of CAMTA3 is promoted via interaction with SR1IP1 (SR1 Interaction Protein 1), a substrate-adaptor for cullin3-based E3 ubiquitin ligase (Zhang et al., 2014). Contrary to above-described understanding of CAMTA3 function as a negative regulator, the enhanced defense phenotype of *camta3* is recently reported to be a form of ETI-like autoimmunity triggered by the activation of two nucleotide-binding domain leucine-rich repeat-containing (NLR) proteins (Lolle et al., 2017). A recent transcriptome analysis further pinpoints CAMTA3 as an early convergence point between PTI and ETI (Jacob et al., 2018). Thus, despite ambiguity about its negative regulator function, CAMTA3 has been independently isolated in a number of disease resistance genetic screens and is clearly important for plant immunity (Du et al., 2009; Jing et al., 2011; Nie et al., 2012).

In contrast to its role as a repressor, CAMTA3 was initially reported to act as transcriptional activator in cold stress response. For instance, expression of the cold-responsive gene *CBF2* (*C-repeat-Binding factor 2*) is positively regulated by CAMTA3 in response to cold stress (Doherty et al., 2009). It was also found to be a transcriptional activator in general stress response and regulation of glucosinolate metabolism (Laluk et al., 2012; Benn et al., 2014). Therefore, CAMTA3 appears to function as a transcriptional repressor for immunity-related genes but acts as a transcriptional activator when bound to promoters of genes involved in cold acclimation or other general stresses. How CAMTA3 switches between activator and repressor functions remains unclear. One possibility is through molecular interactions, either with other proteins or intra-molecularly. A recent study showed that in unstressed plants, CAMTA3 represses transcription of SA pathway genes through an N-terminal repression module (NRM) of CAMTA3; cold stress promotes CaM binding to IQ and/or CaMB domains, resulting in conformation change that interferes with the repressor activity of the NRM (Kim et al., 2017). Additionally, post-translational modifications such as phosphorylation may also be involved (Zhang et al., 2014). Proteomics analysis identified CAMTA3 as a putative MPK3/6 substrate (Hoehenwarter et al., 2012). In addition, in the PhosPhAt database (an Arabidopsis database for experimentally determined phosphorylation sites) (Heazlewood et al., 2008), five phosphopeptides are annotated, which contain typical MAPK phosphorylation sites.

In this work, we investigated CAMTA3 phosphorylation and how this affects its function. We show that CAMTA3 is indeed phosphorylated by flg22-responsive MAPKs, which promotes its destabilization. We could not confirm physical interaction between CAMTA3 and SR1IP1 but show that they have distinct cellular localization and therefore cannot physically interact prior to PTI activation. Finally, we also uncovered that there are additional flg22-responsive kinases that phosphorylate CAMTA3, which may be a CDPK or another kinase downstream of CDPK.

## Results

### CAMTA3 negatively regulates defense-related gene expression

Datamining of expression databases (Hruz et al., 2008) indicated that besides salicylic acid, treatment with PAMPs (e.g. flg22, elf26 or chitin) induces the accumulation of *CAMTA3* transcripts. By RT-qPCR, we confirmed the increase in *CAMTA3* expression in *Arabidopsis thaliana* Col-0 seedlings stimulated with flg22 (Fig. 1A). *CAMTA3* mRNA levels increased rapidly within 30-60 min of flg22 treatment and subsided to near basal levels after ~4 h. Since CAMTA3 is a transcription factor, this increased expression may contribute to the flg22-induced transcriptional response. CAMTA3 is known to regulate expression positively, such as for the cold-responsive gene, *CBF2* (Doherty et al., 2009), or negatively, as was shown for several defense genes, e.g. *EDS1* (Du et al., 2009), *NDR1* and *EIN3* (Nie et al., 2012). Notably, promoters of all of these genes contain CGCG box as *cis*-elements. To investigate the effect of CAMTA3 on PAMP-regulated gene expression, CAMTA3 overexpression (OE) lines were generated. The enhanced basal expression (i.e. in the mock treated tissues) of *CAMTA3* in two independent OE lines was confirmed by RT-qPCR. Flg22 treatment further raised CAMTA3 expression in the OE lines (Fig. 1B), which is presumably the sum of expression arising from the transgene and the inducible expression of the endogenous gene. Next, we analyzed the effect of CAMTA3 on expression of *EDS1*. As expected, *camta3* mutants show high basal *EDS1* transcript levels and CAMTA3 overexpression reduced *EDS1* transcript accumulation. Flg22-induced *EDS1* expression was enhanced in the *camta3* mutants and reciprocally, it was reduced in the two independent OE lines (Fig. 1C). A similar effect was also shown on *NHL10* (Fig. 1D), which has not been reported as a direct CAMTA3 target but includes a potential CAMTA3-binding site in its promoter. These observations are consistent with CAMTA3 functioning as a negative regulator (repressor) during plant immunity response. The PAMP-induced early expression of CAMTA3 may thus potentially serve to shut down expression of downstream target genes.

**Fig 1.**
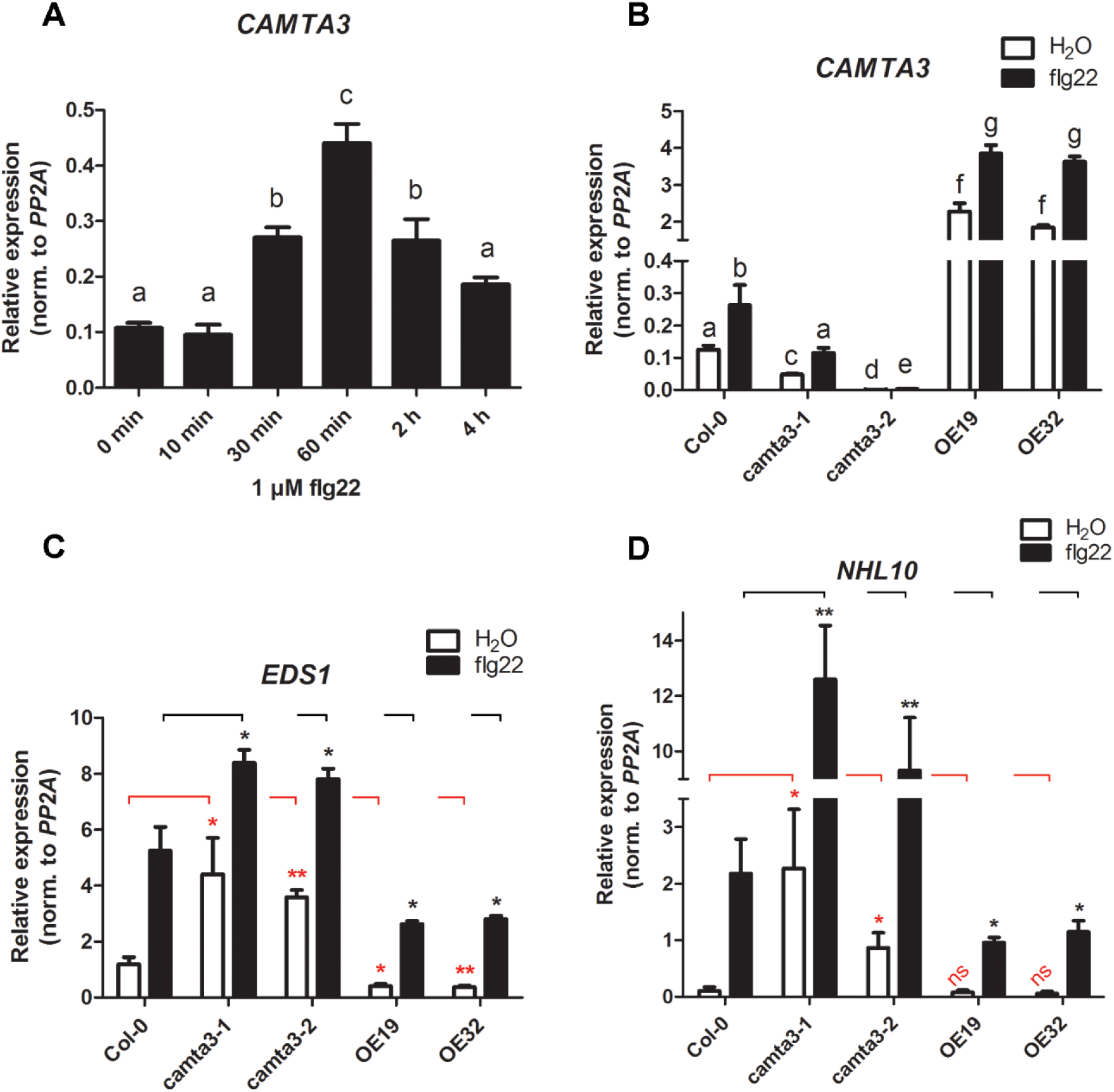
CAMTA3 negatively affects expression of defence-related genes upon flg22 treatment. **A.** Time course of flg22-induced *CAMTA3* expression. Arabidopsis Col-0 seedlings (10-day-old) were elicited with 1 µM flg22, and harvested at the indicated time-points and processed for RT-qPCR. Bars with different letters are significantly different (One-way ANOVA Newman-Keuls multiple comparison test). Error bars = ± SD (n=4). **B.** Analysis of *CAMTA3* transcript levels in mutant and overexpressing (OE) lines upon flg22 or H_2_O treatment. Leaves of 5-week-old plants of the indicated genotypes were infiltrated with H_2_O or 1 µM flg22, harvested after 1 hour and processed for RT-qPCR. OE19 and OE32 are two independent p35S::CAMTA3-YFP overexpressing plants. Bars with different letters are significantly different (One-way ANOVA Newman-Keuls multiple comparison test). Error bars = ± SD (n=6). **C-D.** *EDS1* and *NHL10* expression in the indicated genotype upon flg22 or H_2_O treatment (1 h) were analyzed as described in B. Pairwise t-test (to the corresponding Col-0 controls, as indicated by red or black horizontal brackets) was used to evaluate significance in expression (*p<0.05; **p<0.01, n=3).

### flg22 induces CAMTA3 phosphorylation and destabilization *in vivo*

To investigate the effect of flg22 treatment on the CAMTA3 protein levels, western blot analysis was performed with protein extracted from Arabidopsis protoplasts that transiently express HA-epitope-tagged CAMTA3 under the control of a constitutive CaMV 35S promoter, which thereby uncouple protein accumulation from effects from inducible expression of the native *CAMTA3* promoter. The strong 35S promoter may, however, mask subtle changes in protein levels. To further facilitate visualization of changes in protein turnover, cycloheximide (CHX), a protein translation inhibitor, was included during elicitation. Compared to the water-treated control samples, the CAMTA3 protein level was reduced upon flg22 treatment (in the presence of CHX) (Fig. 2A, upper panel), suggesting that CAMTA3 has a shorter half-life after flg22 treatment. Proteasome-mediated degradation is apparently involved since this destabilization could be blocked by MG115 co-treatment (Fig. 2A, lower panel). Hence, in agreement to repressor functions of CAMTA3, the findings support a model where CAMTA3 destabilization upon PAMP treatment may lead to its removal from promoters and the de-repression of defense gene expression. In the native context, flg22-inducible *CAMTA3* expression (Fig. 1A) may reflect the replacement of the degraded CAMTA3 proteins and hence a negative feedback regulation to restore repression.

**Fig 2.**
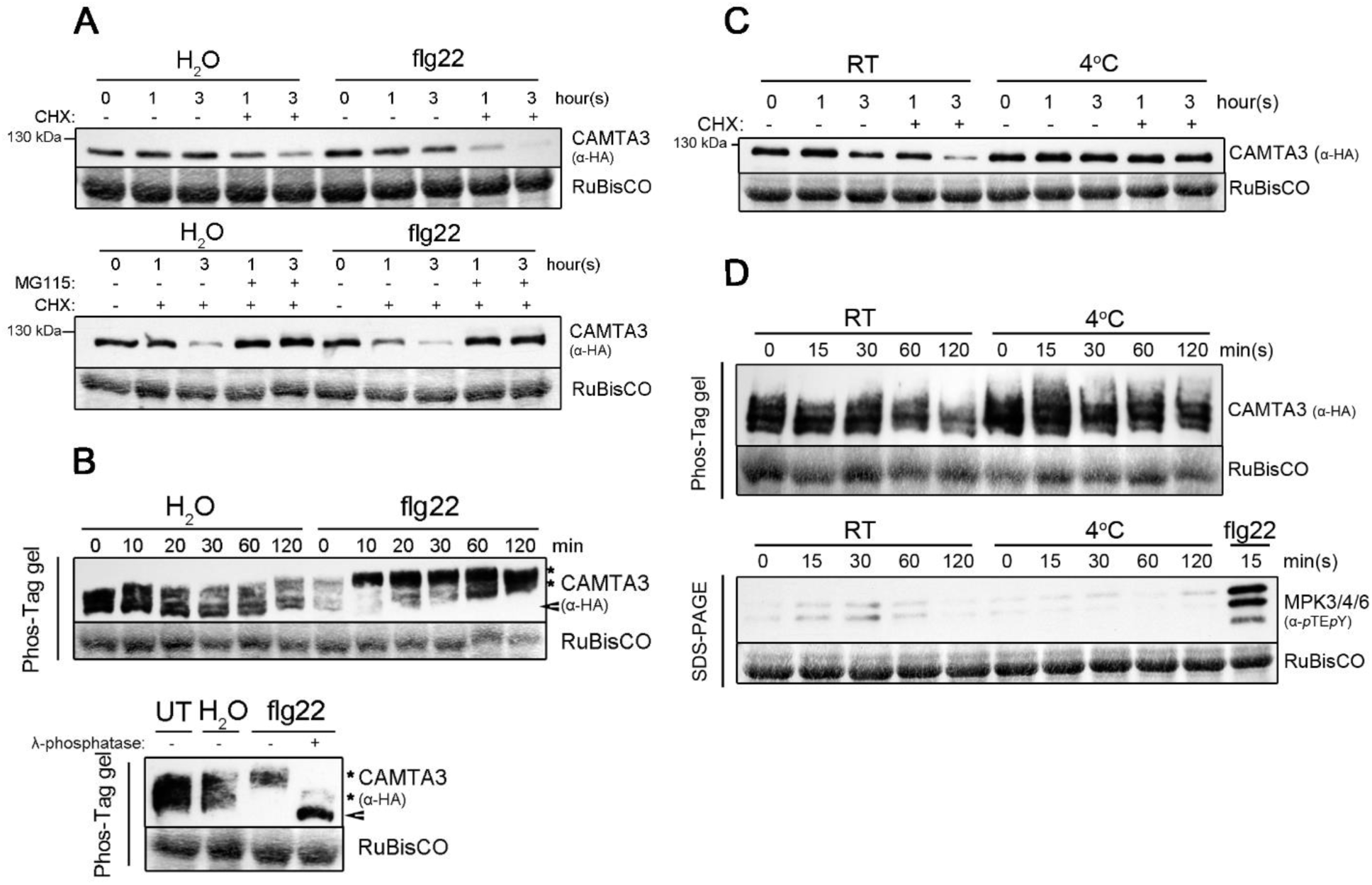
Flg22 induces CAMTA3 phosphorylation and destabilization *in vivo*. **A.** Flg22-induced CAMTA3 protein destabilization is dependent on the proteasome pathway. Upper panel: Protoplasts transfected with plasmids expressing HA-epitope-tagged CAMTA3 were treated with 100 nM flg22 or H_2_O (as control), and proteins were extracted for standard SDS-PAGE and western blot analysis (anti-HA antibody). Where indicated, cycloheximide (CHX, 1 µM) co-treatment was included to block protein translation and facilitate visualization of differential protein turnover. Bottom panel: Protoplasts were pretreated with MG115 (50 µM for 30 min) before elicitation. **B.** PAMP-induced phospho-mobility shift of CAMTA3. Upper panel: Extracted proteins were separated on Manganese(II)-Phos-Tag^TM^-based SDS-PAGE and analyzed by western blot. Bottom panel: Protein extracts were incubated with λ-phosphatase (37°C, 10 min) before Phos-Tag^TM^-based separation and western blotting. Arrowhead marks the “unmodified” CAMTA3 and *marks the various phosphorylated CAMTA3 forms. Amido black staining of the large subunit of RuBisCO was used as loading control for all western blot analysis. **C.** Cold treatment does not reduce CAMTA3 protein stability. Transfected protoplasts were resuspended in media at 4°C or room temperature, harvested at the indicated time points and processed for standard western blotting. **D.** Cold treatment was performed as described in C and the proteins analyzed for phospho-mobility shift of CAMTA3 after separation on Phos-Tag^TM^-based SDS-PAGE. Lower panel: Western blot with phosphorylation specific α-*p*TE*p*Y antibody was used to monitor MAPK activation. Flg22 treatment was used as a positive control for displaying the three PAMP-responsive MAPKs.

Phosphorylation has been speculated to be involved in the destabilization of CAMTA3 after bacterial infection (Zhang et al., 2014). We therefore investigated phosphorylation status upon flg22 elicitation. Here, we employed Manganese (II)-Phos-Tag^TM^-based western blot analysis, where phosphorylated forms show retarded mobility in the gels compared to the non-phosphorylated proteins (Kinoshita et al., 2009). Unlike the well-defined single band in standard PAGE, CAMTA3 from unstressed or H_2_O treated protoplasts already appears as broad smeary bands on Phos-Tag-based western blot, thus suggesting that it exists as several partially phosphorylated forms. By contrast, a mobility shift was seen for CAMTA3 from flg22-treated protoplasts within 10 min of treatment (Fig. 2B, upper panel). λ-phosphatase treatment of the protoplast extracts (Fig. 2B, lower panel) abrogated this mobility shift, suggesting that the smeary bands or reduced mobility bands are due to phosphorylation. Taken together, the Phos-Tag and the phosphatase analyses indicate an *in vivo* flg22-induced CAMTA3 phosphorylation.

By contrast, cold stress (transfer to ice-cold media and placed into a 4°C cold room) did not induce any CAMTA3 degradation within the tested time period and in fact, there appeared to be enhanced protein stability (or less protein degradation) despite termination of protein translation by CHX treatment (Fig. 2C). This is in line with CAMTA3 functioning as a transcriptional activator for cold stress (Doherty et al., 2009). Cold treatment also did not lead to phospho-mobility shift of CAMTA3 in the Phos-Tag analysis (Fig. 2D, upper panel), where MAPKs were correspondingly not activated within the tested time points (Fig. 2D, lower panel). Taken together, unlike PAMP (flg22) elicitation, cold stress does not trigger CAMTA3 phosphorylation and degradation, suggesting differential mechanism exists to determine positive or negative functions of CAMTA3 in these distinct stress context.

### CAMTA3 interacts with and is phosphorylated by PAMP-responsive MAPKs

MAPKs are rapidly activated after flg22 treatment and are thus candidate kinases responsible for the CAMTA3 phosphorylation. The best studied MAPKs are MPK3, MPK4 and MPK6, which are known to be activated by flg22 (Asai et al., 2002). Reconstitution of YFP signals in the bimolecular fluorescence complementation (BiFC) assays suggest that CAMTA3 can potentially interact directly with MPK3, MPK4, and MPK6 (Fig. 3A). By contrast, no YFP signal was seen with an unrelated MAPK (MPK8), suggesting some specificity of the BiFC assay. To investigate whether CAMTA3 is phosphorylated by flg22-responsive MAPKs, an *in vitro* kinase assay was performed with recombinant His-tagged CAMTA3 and GST-tagged MPK3, MPK4 or MPK6. While nearly similar levels of *in vitro* autophosphorylation of the kinases were seen, CAMTA3 was phosphorylated by MPK3 and MPK6, but barely by MPK4 (Fig 3B).

**Fig 3.**
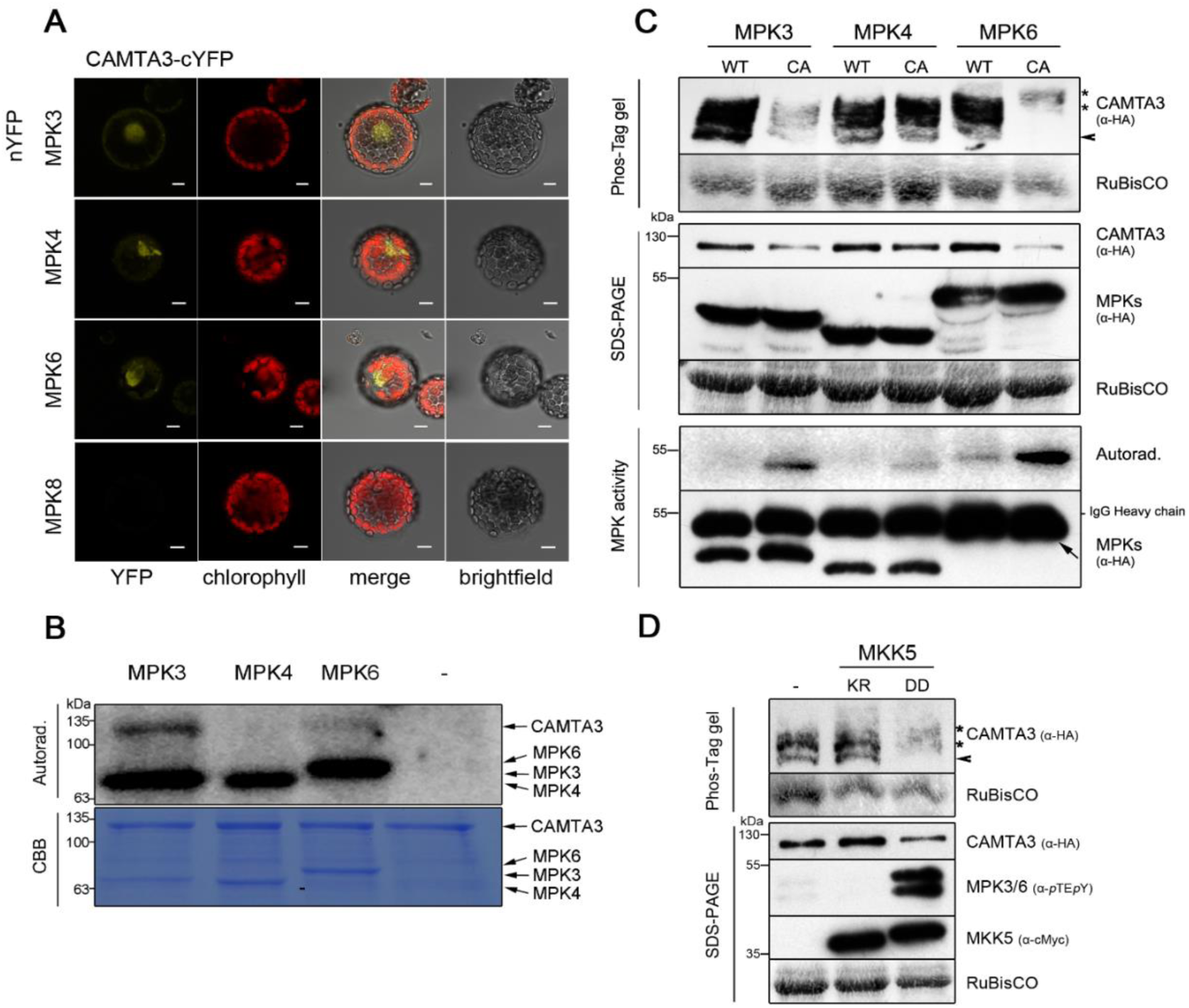
CAMTA3 interacts with and is phosphorylated by PAMP-responsive MAPKs. **A.** CAMTA3 interacts with stress-activated MAPKs (MPK3, MPK4, and MPK6). Bimolecular fluorescence complementation (BiFC) assay was performed in protoplasts co-expressing CAMTA3 (fused with a C-terminal fragment of YFP, cYFP) and MPK3 or MPK4 or MPK6 (fused with an N-terminal fragment of YFP, nYFP). MPK8, a non PAMP-responsive MAPK, was used as a negative control. Interaction is visualized as reconstituted YFP signals. Scale bar = 10 µm. Western blot validation detecting intact fusion proteins is shown in Fig. S6A. **B.** *In vitro* kinase assay of recombinant CAMTA3 by PAMP-responsive MAPKs. Recombinant His-tagged CAMTA3 (as a substrate) was incubated with the indicated GST-tagged MAPKs in the presence of ^32^P-labelled ATP (for 30 min at 37°C). After SDS-PAGE separation, phosphorylated proteins were visualized by autoradiography. Lower panel is the corresponding coomassie blue staining (CBB). **C-D.** CAMTA3 is phosphorylated mainly by MPK3 and MPK6 *in vivo*. **C.** Transient co-expression of constitutively-active (CA) MPK3 and MPK6, but not CA-MPK4 or wild-type (WT) MPKs, led to increased phosphorylation and destabilization of CAMTA3. Protoplasts were transfected with the indicated constructs and the extracted proteins were analyzed by western blotting after separation on Phos-Tag gel or conventional SDS-PAGE to monitor phospho-mobility shifts (upper panel) or protein stability (middle panel), respectively. In the bottom panel, CA-MAPKs and WT-MAPKs transiently expressed in protoplasts were immunoprecipitated using anti-HA antibody and their auto-phosphorylation (in the presence of ^32^P-labelled ATP) used to compare their kinase activities. Western blot was used to estimate expression levels of the MAPKs. Note that the MPK6 immunological signals (see arrows) partially overlaps with the IgG heavy chain signals. **D.** Co-expression of a constitutively active MKK5 (DD), but not its kinase-dead variant (KR), led to phospho-mobility shift and reduced protein stability of CAMTA3. CAMTA3 was transiently expressed in protoplasts together with MKK5^DD^, which activates MPK3/6 activation. (The kinase-inactive variant, MKK5^KR^ was used as as control). After an overnight incubation for protein expression, Phos-Tag based or conventional SDS-PAGE based western blots with the indicated bodies were performed as described above. Expression of CAMTA3 was monitored with anti-HA antibody, and specific MPK3 and MPK6 phosphorylation by MKK5^DD^ with α-*p*TE*p*Y antibody, and MKK5 (DD and KR) with anti-cMyc antibody. Arrowhead= “unmodified” CAMTA3; *= phosphorylated CAMTA3 forms.

To verify *in vivo* phosphorylation, constitutively active MAPKs (CA-MAPKs) (Berriri et al., 2012; Genot et al., 2017) were transiently co-expressed with CAMTA3 in protoplasts. Increased kinase activities of these CA-MAPKs compared to the native kinases were demonstrated by the higher auto-phosphorylation of the immunoprecipitated MAPKs. *In vivo* CAMTA3 phosphorylation and any effect on the protein levels were then monitored by Phos-Tag or standard western blot, respectively. CAMTA3 was phospho-shifted when co-expressed with CA-MPK6 and accumulated to lower levels. For CA-MPK3 co-expression, some phosphorylation (but weaker compared to MPK6) was observed but a strong CAMTA3 destabilization was seen. Consistent with *in vitro* kinase assay, CA-MPK4 co-expression did not induce CAMTA3 phospho-shift (Fig. 3C). Thus, MPK4 is probably not involved in inducing *in vivo* phosphorylation and destabilization of CAMTA3. However, some caveats to the above interpretations include: (1) the CA-MPK4 displayed lower autophosphorylation activity than the other two kinases; (2) it cannot be excluded that substrate specificities or activities of these mutated MAPKs are comparable to the MAPKs natively activated through upstream MAPK kinases.

To validate our results, we transiently expressed in protoplasts a constitutively active MKK5 (MKK5^DD^) to specifically activate endogenous MPK3 and MPK6 but not MPK4 (Lee et al., 2004; Lassowskat et al., 2014). In this case, MPK3/6 are activated “naturally” through phosphorylation of its kinase activation loop and not through mutation. Similar to observations above for CA-MPK3 and CA-MPK6, CAMTA3 protein level was reduced through MKK5^DD^ co-expression. The phosphorylated CAMTA3 proteins are difficult to visualize in the Phos-Tag gels since the strongly reduced protein levels are further separated into smeary, barely detectable, bands. However, the slower mobility of these fuzzy bands is in agreement with MPK3/6-mediated phosphorylation. Control transfection without any MKKs or co-transfection with a kinase-inactive MKK5 (MKK5^KR^) did not show these effects (Fig. 3D). Hence, the results above suggest that two out of three known flg22-responsive MAPKs phosphorylate CAMTA3, and *in vivo* flg22-induced CAMTA3 destabilization may be triggered by phosphorylation via MPK3 and MPK6.

### CAMTA3 is phosphorylated by PAMP-responsive MPK3/6 at multiple sites, which contribute to CAMTA3 protein destabilization

After confirming *in vitro* and *in vivo* CAMTA3 phosphorylation by MPK3 and MPK6, we proceeded to determine the phospho-sites in CAMTA3. CAMTA3 contains 11 potential MAPKs phosphorylation sites, which are serine or threonine followed by a proline (S/TP) (Fig. 4A). Both *in vitro* kinase assay (with non-radioactive ATP) and infiltrating flg22 to CAMTA3-overexpression lines were used for collecting phosphorylated CAMTA3 protein samples. MPK3-or MPK6-phosphorylated CAMTA3 bands, or flg22-induced phosphorylated CAMTA3 proteins immunoprecipitated from overexpression plant extracts, were excised after SDS-PAGE separation. After tryptic digestion and phospho-peptide enrichment, samples were subjected to mass spectrometry (MS) analysis (MS2 spectra mapping the respective phospho-sites are shown in Fig. S1). Three phospho-peptides with phosphorylated S272, S454, or S780 were detected (Fig. 4A, marked in red). PhosPhAt 4.0 database, an *Arabidopsis thaliana* phosphorylation site database incorporating experimental phosphoproteomic data (Heazlewood et al., 2008), annotates two further phosphorylation sites at S8 (Hoehenwarter et al., 2012) and S587 in CAMTA3 (Fig. 4A, marked in blue). Based on these data, we generated phospho-mutants: CAMTA3-mutP1 and -mutP2, with respectively three or five sites mutated to non-phosphorylatable residues (Fig. 4A). However, *in vitro* MPK3- or MPK6-mediated phosphorylation of the CAMTA3-mutP1 and -mutP2 variants was not reduced compared to CAMTA3-WT (Fig. 4B), suggesting the presence of additional MAPK-targeted sites.

**Fig 4.**
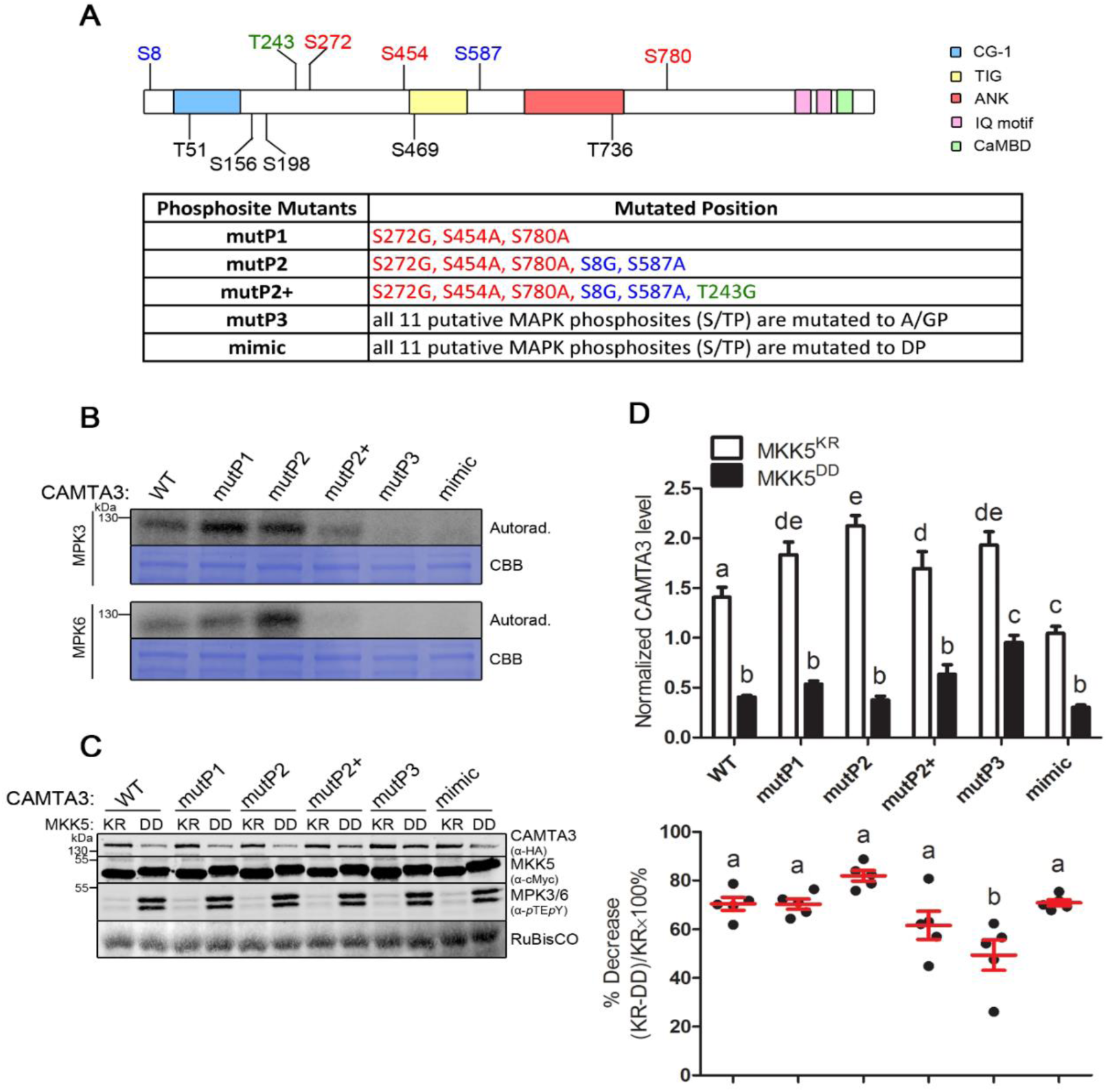
CAMTA3 is phosphorylated by MPK3/6 at multiple sites, which contribute to CAMTA3 destabilization. **A.** Schematic depiction of conserved domains within CAMTA3. (CG-1= CG-box binding domain; TIG= immunoglobulin-fold domain (putative unspecific DNA binding); ANK= Ankyrin repeat; IQ motif/CaMBD= Calmodulin binding motifs). The 11 possible MAPK phospho-sites are indicated marked (Color code: Red= phosphosite identified by mass spec after tryptic digest, blue= phosphosite annotated in PhosPHAT; green= novel phosphosite identified after optimizing digestion). Table summarizes the different phosphosite mutations generated in this study. (Note substitution to alanine was used to generate non-phosphorytable residues in all cases except at S8, S198, S272, T243, S469 and T736, where glycine was used due to the convenience of the cloning procedure. See methods for details). **B.** *In vitro* kinase assay with WT and phosphosite mutants of CAMTA3. The indicated CAMTA3 variants were incubated active MPK3 or MPK6 and ^32^P-labelled ATP. Shown are the autoradiograph of phosphorylated CAMTA3 bands and the corresponding coomassie blue staining (CBB). **C.** MPK3/6-mediated phosphorylation of CAMTA3 promotes degradation. CAMTA3 variants were transiently co-expressed in protoplasts with constitutively active MKK5 (DD) or a kinase-deficient version (KR) as negative control. Extracted proteins were analyzed by western blotting to detect the abundance of CAMTA3 (α-HA), MKK5 (α-cMyc) and the activated MPK3/6 (α-*p*TE*p*Y). Amido black staining of the RuBisCo band served as a loading control. This experiment was repeated eight times and a representative blot shown. **D.** Quantitative western blot estimation of CAMTA3 abundance. After transient expression and western blotting as described in C, CAMTA3 abundance was quantified as density of protein bands of five replicates (each consisting of independent protoplast samples). The absolute values from individual experiment were normalized by the median of each dataset and analyzed using one-way ANOVA (*p* < 0.05). To take into account the different basal levels of the different protein variants, percentage decrease comparing between KR and DD was additionally calculated (within each experiment), using the formula (KR-DD)/KR×100. Error bars = SEM. Statistically distinct groups after 1-way ANOVA are marked with different alphabets (both upper and lower panels).

To increase MS sequence coverage of CAMTA3 we repeated the experiment but digested the protein with both trypsin and endoproteinase Glu-C prior to MS analysis. This resulted in more than 60% sequence coverage that encompass most of the SP/TP motifs, leading to the identification of phosphorylation at T243, S587 and S780 (Fig. S1). Notably, T243 (Fig. 4A, marked in green) is a novel phospho-site, never detected in any previous reports. When the phospho-mutant CAMTA3-mutP2+ (containing phospho-site mutations at S8, T243, S272, S454, S587 and S780) was tested, *in vitro* MPK3- or MPK6-mediated phosphorylation was strongly reduced compared to CAMTA3-WT (Fig. 4B), so that these six sites are likely the major MPK3/6-targeted sites. We also generated CAMTA3-mutP3 (as a phospho-null for all 11 potential phospho-sites) and the corresponding phospho-mimetic mutant (CAMTA3-mimic), in which all 11 potential sites (S/T) were substituted by aspartic acids (D) (Fig. 4A). Phosphorylation of these variants was completely lost (Fig. 4B), which also proved that MPK3/6 did not unspecifically phosphorylate other residues in CAMTA3.

To investigate the role of phosphorylation in protein destabilization, we compare the stability of the different CAMTA3 phosphosite variants. Co-expression of MKK5^DD^ (or the kinase-inactive MKK5^KR^ variant as a control) was employed to investigate the actions of MPK3/6. A quantitative western blot system (Li-COR Odyssey® CLx multiplex imaging system) was used to determine the protein levels in five replicates (consisting of independent protoplast samples). Fig. 4C shows a representative western blot, and the bar chart in Fig. 4D (upper panel) shows the quantification of the protein levels. Reduction of CAMTA3-WT protein level was seen when MAPK activation was induced by MKK5^DD^ co-expression (Fig. 4C); statistical analysis shows that this is also true for all the mutated CAMTA3 (Fig. 4D). Since MKK5^DD^ may potentially activate the proteasome pathway (Lee et al., 2015), subtle differences in protein stability may be partially masked. However, note that basal expression of all the phospho-site mutants are higher levels than the CAMTA3-WT while the opposite is true for the CAMTA3-mimic (Fig 4D). Thus, the MAPK-targeted phospho-sites may be crucial for maintaining protein stability.

As seen in the representative western blot (Fig. 4C), the mutP2+ and mutP3 phospho-mutants appear slightly more stable than the CAMTA3-WT protein after MPK3/6 activation, which is substantiated by statistical analysis for the mutP3 variant. Although interpretation for mutP2+ lacked statistical robustness, this may be a type II (false negative) statistical error because we do see this qualitative difference in most of the independent replicates. As seen in the recalculation of the percent decrease (Fig. 4D, lower panel), one (out of the five) datapoint may be an outlier and the average percent decrease of mutP2+ is intermediate between WT and mutP3. Taken together, these results indicate that CAMTA3 destabilization requires phosphorylation at multiple MPK3/6-targeted sites, including the six major phospho-sites analyzed in the mutP2+ variant.

The potential mechanism behind CAMTA3 destabilization through phosphorylation is still unknown. A previous study proposed that the CAMTA3-interacting SR1IP1 recruits CAMTA3 for ubiquitination and degradation by the 26S proteasome (Zhang et al., 2014). To test the involvement of SR1IP1, we aimed to check if phosphorylation affected SR1IP1-CAMTA3 interaction. We performed several assays including yeast two-hybrid, bimolecular fluorescence complementation (BiFC) and split-assays in transfected Arabidopsis protoplasts (Fig. S2), but none of the assays supported interaction between SR1IP1 and CAMTA3.

### MAPK-induced phosphorylation triggers CAMTA3 subcellular relocalization

To clarify the discrepancy to the literature, we looked at the cellular distribution of the two proteins. CAMTA3 is a transcription factor and mainly localized in the nucleus (Yang and Poovaiah, 2002), which we could also confirm in unstressed protoplasts using a CAMTA3-YFP fusion construct (Fig. 5A, upper panel). However, SR1IP1-CFP signals are in the periphery (presumably plasma membrane) of the protoplasts (Fig. 5A, lower panel). Hence, the different localization might explain why we could not detect any interaction between SR1IP1 and CAMTA3 in any of the plant cell-based assays. For yeast two-hybrid assay, we employed a GAL4-based system while the reported SR1IP1-CAMTA3 interaction is based on the *pSOS* system (Zhang et al., 2014), which relies on myristylation of the bait proteins. Thus, the recruitment of the reporter to the plasma membrane may permit interaction. Nevertheless, the distinct localization of CAMTA3 and SR1IP1 suggests that they cannot physically interact unless there is a re-localization of either proteins from the nucleus or the plasma membrane, respectively.

**Fig 5.**
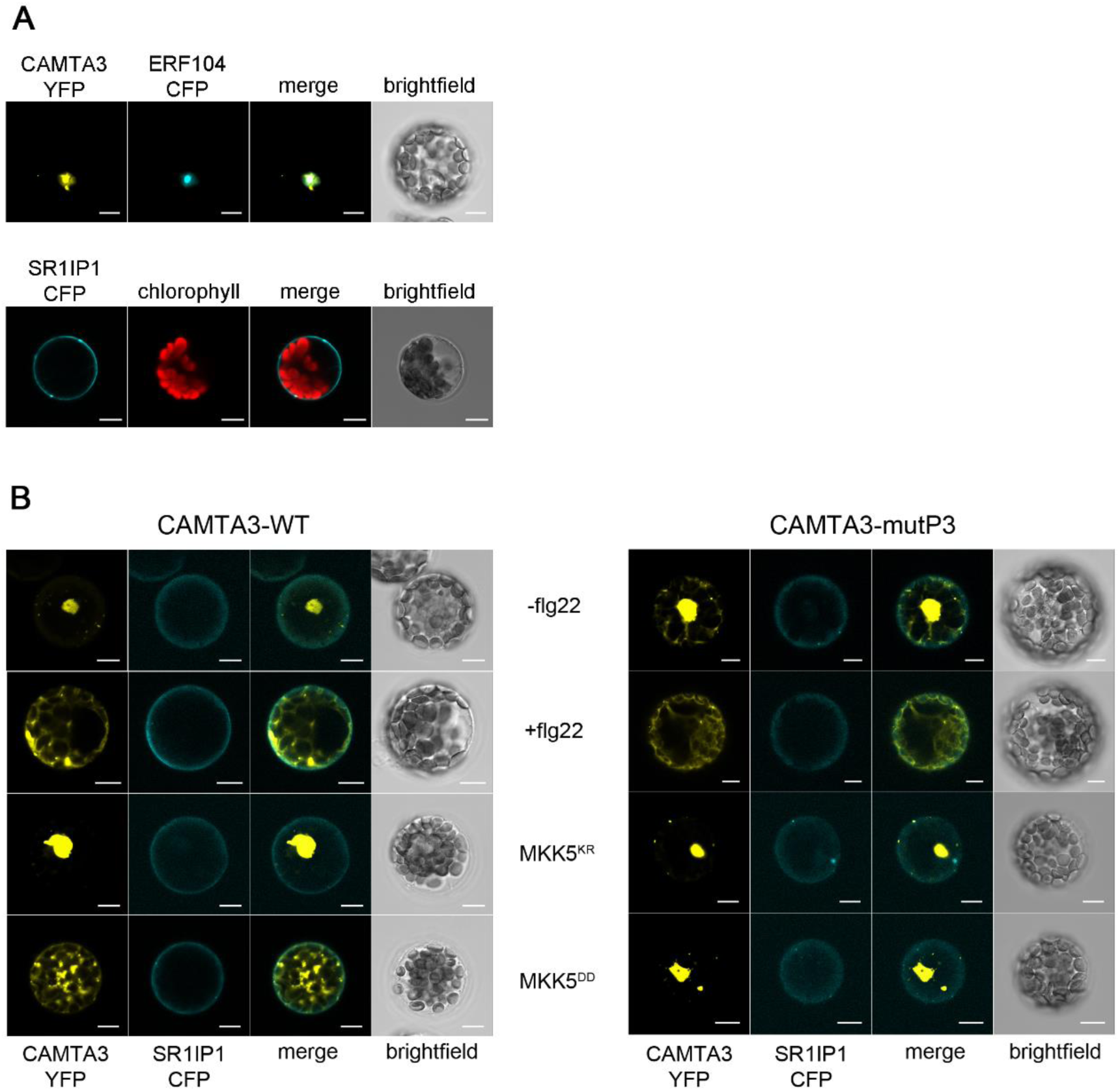
Flg22 and MPK3/6-mediated phosphorylation induce nuclear export of CAMTA3. **A.** CAMTA3 and SR1IP1 are localized in different cell compartments. Top panel: CAMTA3-YFP was co-expressed with ERF104-CFP (used as a nuclear marker) in protoplasts. Bottom panel: SR1IP1-CFP was transiently expressed in protoplasts. After protein accumulating overnight, the protoplasts were observed by confocal microscopy. Position of chloroplasts are visualized through the chlorophyll autofluorescence. Scale bar = 10 µm. **B.** Protoplasts co-expressing CAMTA3-YFP and SR1IP1-CFP were treated with flg22 (100 nM, for 2h) or water as control (-flg22) and observed by confocal microscopy. Bottom row shows additional co-transfection with MKK5^DD^ (to activate MPK3 and MPK6). The inactive MKK5^KR^ serves as a negative control. Protoplasts were observed by confocal microscopy after an overnight incubation. The right panel shows the same experiment set up except the phospho-null CAMTA3-mutP3 was used in place of the WT CAMTA3. Scale bar = 10 µm. Western blot detection of intact proteins is shown in Fig. S6B.

Interestingly, the CAMTA3-YFP signal relocated from nucleus to the cytoplasm after flg22 elicitation, and multiple cytoplasmic aggregates were often observed (Fig. 5B; see additionally the time course experiment in Fig S3A). The aggregates are not cleaved YFP fragments since western blot analysis revealed intact CAMTA3-YFP protein bands (Fig. S3B). By contrast, SR1IP1 localization was not visibly affected by flg22 treatment. Although the CAMTA3 relocalization into the cytoplasm may potentially allow interaction with SR1IP1 and indeed, there seems to be partial co-localization between CAMTA3-YFP and SR1IP1-CFP after flg22 treatment (Fig. 5B), we never obtain any BiFC or split-LUC signals between CAMTA3 and SR1IP1 even upon flg22 elicitation (Fig. S2). Nonetheless, we cannot exclude that CAMTA3-SR1IP1 does interact upon flg22 elicitation but is degraded too rapidly for detection of the complex.

Notably, activation of MPK3/6 (through MKK5^DD^ co-expression) also induced the CAMTA3 relocalization to the cytoplasm (Fig. 5B, left lower panels), thus phenocopying the effects of flg22 elicitation. The non-phosphorylatable CAMTA3-mutP3 remained nuclear-localized with the MKK5^DD^ co-expression (Fig. 5B, right lower panels), demonstrating that the mutated phospho-sites are crucial for the relocalization and that CAMTA3 nuclear export can be triggered by its MPK3/6-mediated phosphorylation. On the other hand, these phospho-sites also appear to be irrelevant for the flg22 response since the CAMTA3-mutP3 mutant still re-localized to the cytoplasm upon flg22 treatment (Fig. 5B, right panel). In summary, the MAPK-targeted phospho-sites are dispensable for the subcellular relocalization of CAMTA3 induced by flg22 treatment but are essential for the MPK3/6-induced pathway. Thus, in the case of flg22 elicitation, there are alternative (non-MPK3/6) pathways that induce CAMTA3 relocalization.

### Additional kinases may be involved in flg22-mediated phosphorylation of CAMTA3

During the Phos-Tag-based analysis of CAMTA3 phospho-status, we noticed that flg22 treatment can induce additional mobility shift to the phosphoshift already triggered by MKK5^DD^ co-expression (i.e. *in vivo* MPK3/6 activation) (Fig. 6A, left panel). This strong mobility shift was also seen for the mutP3 variant, which showed no difference in the mobility shift between co-expressing MKK5^KR^ and MKK5^DD^ (Fig. 6A, right panel). Taken with the observations of CAMTA3 relocalization above, there are other flg22-responsive kinases (besides MPK3/6) that phosphorylate CAMTA3. The analysis with the mutP3 variant that lacks all MAPK-targeted sites also excludes MPK4 or other MAPKs known to be weakly activated by flg22 (e.g. MPK11 (Bethke et al., 2012; Nitta et al., 2014)).

**Fig 6.**
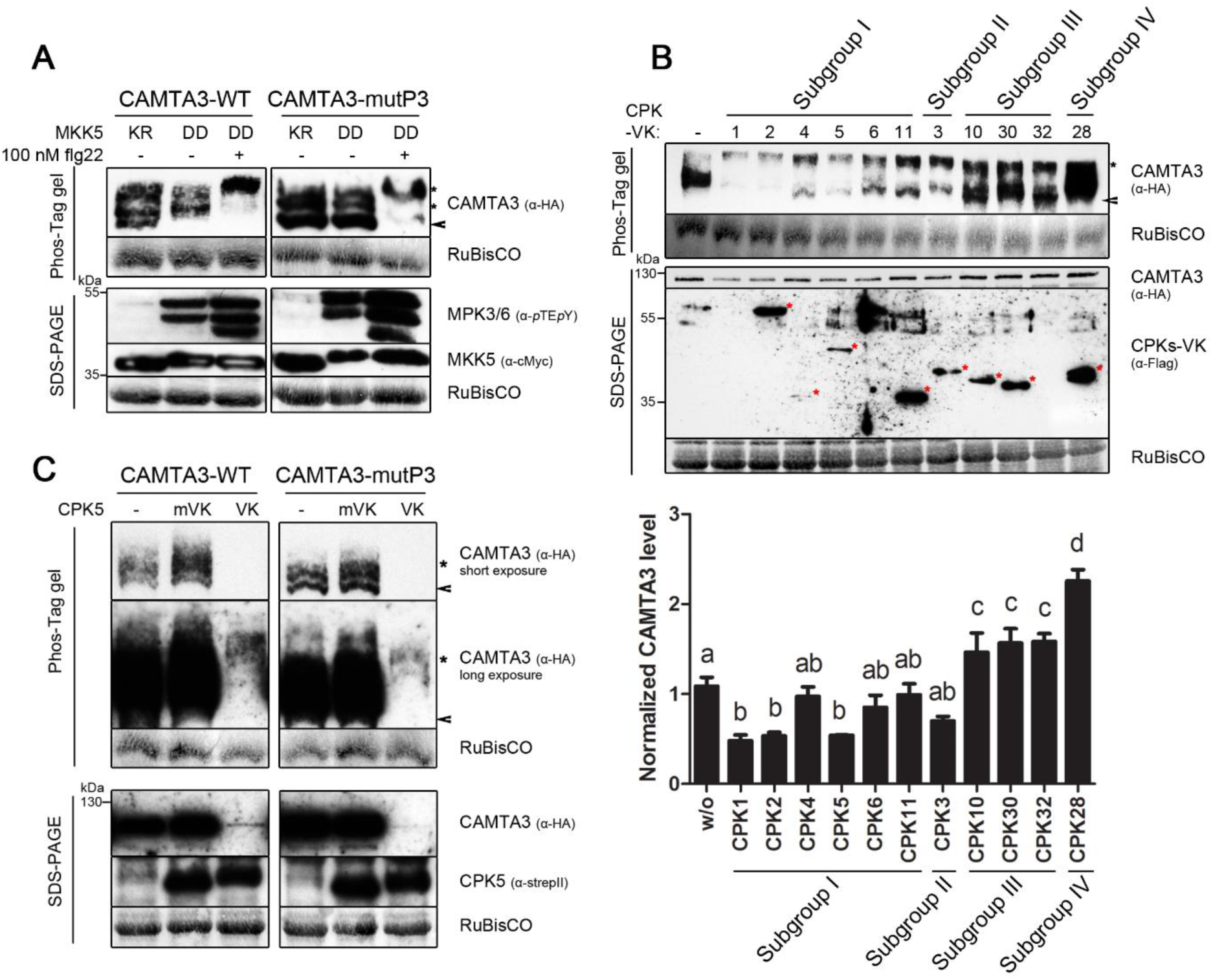
CPK5 may also be involved in flg22-induced phosphorylation of CAMTA3. **A.** Additional kinases are involved in flg22-induced phosphorylation of CAMTA3. Arabidopsis protoplasts were transfected to co-express CAMTA3-WT (or CAMTA3-mutP3) with constitutively active MKK5^DD^ or its kinase-dead variant (KR), respectively. One batch of the protoplasts transfected with MKK5^DD^ was additionally treated with 100 nM flg22 for 10 min. The extracted proteins were analyzed by western blot after separation on Phos-Tag gel to visualize phospho-mobility shift (upper panel) or conventional SDS-PAGE for MKK5 expression and MAPK activation (lower panel). **B.** CDPKs, mainly from subfamily I, induce CAMTA3 phosphorylation and destabilization. CAMTA3 was co-expressed with the indicated Arabidopsis CDPKs, as its constitutively-active variants (designated as VK). Arrowhead= “unmodified” CAMTA3; *= phosphorylated CAMTA3 forms. Middle panel: Red asterisks mark the expected positions of the CDPKs in the anti-Flag western blot. (Note some CDPKs are not detectable despite promoting CAMTA3 destabilization.) Lower panel is the quantitative western blot analysis of CAMTA3 levels after coexpression with the indicated CDPKs (n = 3 independent replicates). Statistical distinct groups (one-way ANOVA, *p* < 0.05) are marked with distinct alphabets. **C.** CPK5-induced destabilization of CAMTA3 is independent of the MAPK-targeted phosphosites. Co-transfection of a constitutively active CPK5 (VK), but not a kinase-deficient variant CPK5m-VK, led to a strong phospho-mobility shift (upper panel) and reduced protein stability (lower panel) of both CAMTA3-WT and mutP3. Note the longer exposure of the western blot to see the phospho-mobility shift.

Besides MAPK cascades, several calcium-dependent protein kinases (CDPKs/CPKs) are rapidly activated after exposure to PAMPs or pathogens (Ludwig et al., 2004). In Arabidopsis, CDPKs are encoded by a large gene family of 34 members and classified into four subfamilies (Boudsocq et al., 2010). Selected members from all four CDPK subfamilies were tested if they are involved in phosphorylation of CAMTA3. Deleting the c-terminal Ca^2+^ regulatory and auto-inhibitory domains of CDPKs creates active kinases. When such constitutively-active CDPKs (designated as VK variants) were transiently co-expressed with CAMTA3 in the protoplasts, most of them induce a phospho-shift of CAMTA3 and, in part, a change in CAMTA3 protein levels (Fig. 6B). According to the quantitative western blot and statistical analysis, CAMTA3 protein stability was significantly reduced by co-expression of CPK1, CPK2, or CPK5, while the tested members from subfamily III and IV did not reduce but increased CAMTA3 levels significantly (Fig. 6B, lower panel). Note that although through overexpression, many of the constitutively-active CDPKs can biochemically modulate CAMTA3 stability, most of them are not known to be PAMP-activated. Among the four Arabidopsis flg22-responsive CPKs (CPK4, 5, 6, and 11) (Boudsocq et al., 2010), only CPK5 highly phosphorylated and destabilized CAMTA3. This is in line with CPK5 being a key calcium-dependent regulator of innate immune responses in plants (Dubiella et al., 2013). Thus, CPK5 is most likely the relevant flg22-responsive kinase that induce MAPK-independent CAMTA3 phosphorylation.

Thus, we tested the role of CPK5 by transiently expressing CAMTA3-WT (or mutP3) together with CPK5-VK or a kinase-deficient version (CPK5m-VK) into protoplasts. CPK5-VK expression led to a strong destabilization of CAMTA3. Extended exposure of the western blots was necessary to visualize the low levels of CAMTA3-WT or mutP3. Importantly, the strong phospho-shift of CAMTA3 induced by CPK5-VK is comparable to the shift induced by flg22 stimulation (Fig. 6C). Since this phospho-shift can be seen in both WT and mutP3 versions of CAMTA3, there is phosphorylation at non-MAPK targeted sites through CPK5-VK. This further supports CPK5 as the unknown flg22-responsive kinase(s) that could target CAMTA3.

### CAMTA3 is not a direct target of CPK5

To investigate whether CAMTA3 may be a target for CPK5, CAMTA3 interaction with CPK5 was tested. BiFC assay performed in protoplasts showed reconstituted YFP fluorescence, suggesting that CAMTA3 can indeed interact directly with full length CPK5. A kinase-deficient version CPK5m was shown to interact with CAMTA3 as well (Fig. 7A, upper panel), which indicates that the interaction between CAMTA3 and CPK5 is not dependent on its kinase activity. BiFC signals co-localized with a nuclear marker protein ERF104 (Bethke et al., 2009), suggesting the interaction occurs mainly in the nucleus (Fig. 7A, bottom panel). All these results indicate that CAMTA3 can interact with CPK5 in the nucleus *in vivo*.

**Fig 7.**
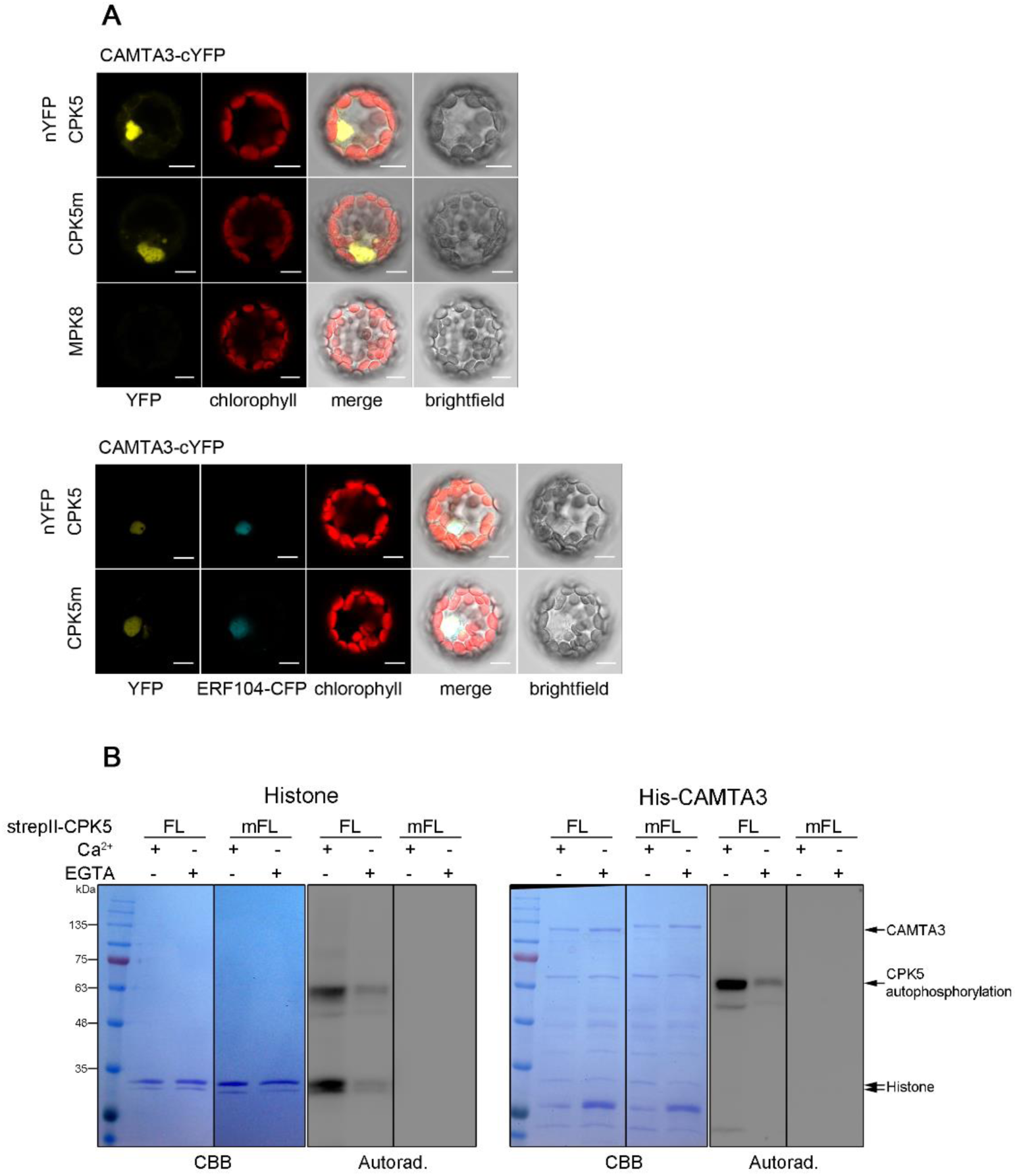
CPK5 does not directly phosphorylate CAMTA3. **A.** CAMTA3 can interact with CPK5 in bimolecular fluorescence complementation (BiFC) assays. Protoplasts were transfected with the indicated combinations of cYFP and nYFP fusions and observed under a confocal fluorescence microscope. MPK8-nYFP fusion was used as a negative control (upper panel). A nuclear marker, ERF104-CFP fusion, was co-expressed to determine localization of the BiFC signals (bottom panel). Scale bar = 10 µm. Western blot validation of intact proteins is shown in Fig. S6C. **B.** CAMTA3 is not phosphorylated by CPK5 *in vitro*. StrepII-tagged full-length CPK5 (CPK5-FL) and a kinase-dead variant (CPK5m-FL) were immunoprecipitated from transfected Arabidopsis protoplasts and incubated (in the presence of 0.5 µg calmodulin and ^32^P-labelled ATP) with either Histone (left panel) or recombinant purified His-tagged CAMTA3 (right panel) as substrates. To show Ca^2+^ dependency of the reaction, EGTA was used to chelate Ca^2+^. Proteins were separated by SDS-PAGE, and phosphorylation was visualized by autoradiography.

After successfully detecting the interaction between CAMTA3 and CPK5, an *in vitro* kinase assay was performed to further investigate whether CPK5 phosphorylates CAMTA3 directly. StrepII-tagged full length CPK5 (CPK5-FL) and a kinase-deficient version, CPK5m-FL, were transiently overexpressed and immunoprecipitated from transfected protoplasts, and purified His-tagged recombinant CAMTA3 was used as a substrate. In the presence of ^32^P-ATP and Ca^2+^, auto-phosphorylation of CPK5-FL and trans-phosphorylation of a generic substrate (histone) were observed (Fig. 7B, left panel). However, no trans-phosphorylation of CAMTA3 was detectable (Fig. 7B, right panel). Calmodulin was additionally included in the reaction because of the role of Ca^2+^/CaM-binding for CAMTA3 functions (Du et al., 2009) but no CAMTA3 phosphorylation was observed CaM was excluded or when constitutively-active CPK5-VK recombinant protein was used (Fig. S5). Taken together, despite an *in vivo* CPK5-induced phospho-shift, CAMTA3 is probably not phosphorylated by CPK5 directly.

## Discussion

CAMTA3 has been reported to act as a transcriptional activator as well as a repressor. The results presented here support the latter, i.e. CAMTA3 is a negative regulator of defence genes, where CAMTA3 (overexpression) suppresses both the basal and flg22-induced expression of defence genes (Fig 1C,D). Furthermore, upon flg22 perception, activated MPK3 and MPK6 can phosphorylate CAMTA3 directly, which promotes CAMTA3 degradation via a proteasome-dependent pathway and also the relocalization of CAMTA3 from the nucleus to cytoplasm. These two processes can thus lead to de-repression of downstream target genes. The transcriptional upregulation of *CAMTA3* by flg22 may thus represent a negative feedback mechanism to replace the degraded CAMTA3 proteins and shut down defence gene expression.

Besides MAPKs, additional flg22-responsive kinase(s) can induce CAMTA3 phosphorylation that strongly promotes its degradation. This is most likely an unknown kinase downstream of CPK5; although we cannot exclude that CPK5 can directly phosphorylate CAMTA3 under optimal conditions (such as the presence of additional co-factors). Taken together, our findings suggest that CAMTA3 may be a converging point of two major flg22-activated phosphorylation pathways, namely MAPK cascades and CDPKs. Boudsocq et al. (2010) showed that activation of CDPKs and MAPK cascades act independently, which is demonstrated by unaltered flg22-induced MAPK activation profiles in the *cpk5 cpk6 cpk11* triple mutant or in plants expressing constitutively-active CDPKs. On the other hand, MAPKs and CDPKs can also function synergistically to regulate the expression of some PTI genes (e.g. *NHL10*, *CYP82C2* and *PER4*) (Boudsocq and Sheen, 2013). Whether CAMTA3 plays a role in this signalling convergence remains to be determined. The typical phosphorylation motifs in MAPK and CDPK substrates are different. The minimal MAPK targeted site is S/T-P, while the predicted CDPK target motif is φ_−1_-[ST]_0_-φ_+1_-X-Basic_+3_-Basic_+4_ (φ is a hydrophobic residue) (Sebastià et al., 2004). Different phosphorylation motifs may, of course, suggest that the MAPKs and CDPKs regulate PTI by targeting different substrates. However, the same target protein might also be phosphorylated by both kinases at different sites. For instance, ACS2, the key enzyme of ethylene biosynthesis, is known to be MPK3 and MPK6 substrate (Liu and Zhang, 2004; Han et al., 2010) but it also bears a CDPK phosphorylation site (Lyzenga and Stone, 2012). By analogy, the tomato ACS2 homolog is phosphorylated by tomato CPK2 and MAPK at different sites in response to wound signalling (Kamiyoshihara et al., 2010). Compared to MAPKs, fewer pathogen-responsive CDPK substrates have been identified. These include the Arabidopsis RBOHD, a substrate of CPK5 after PAMP stimulation (Dubiella et al., 2013), or WRKY8, 28 and 48 transcription factors, which are substrates of CPK4, 5, 6 and 11 (Gao et al., 2013). However, none of these have been shown to be also MAPK targets. To understand the possible crosstalk between MAPK- and CDPK-mediated phosphorylation on CAMTA3, it would be crucial to first identify the putative “intermediate” kinase acting downstream of CPK5 that phosphorylate CAMTA3 (or the conditions needed for CPK5 to directly phosphorylate CAMTA3). Since CAMTA3 physically interacted *in vivo* with CPK5 in the nucleus, both may be needed to identify the required associated component(s). Alternatively, some form of CAMTA3 post-translational modifications may be required before direct phosphorylation by CPK5. However, this is unlikely to be through MAPK phosphorylation since CPK5 or flg22 treatment can both still induce a phospho-shift of the CAMTA3-P3 mutant (where all 11 potential MAPK phosphorylation sites are mutated).

In agreement to pathogen-induced CAMTA3 degradation, we show that flg22-induced CAMTA3 phosphorylation leads to its *in vivo* destabilization and propose this may de-repress expression of downstream target genes. Equally possible would be a direct phospho-dependent reduction in DNA binding properties of CAMTA3. If phosphorylation is involved, only allosteric effects can be expected since the key MAPK-targeted phosphosites are not within the DNA-binding domain (see Fig. 4A). Unfortunately, we were unable to obtain convincing EMSA (electrophoretic mobility shift assay) data proving specific CAMTA3 binding to “CGCG”-core containing DNA probes (Fig S4). All published EMSA results employed truncated CAMTA3 comprising only the DNA-binding domain (Du et al., 2009; Nie et al., 2012) while we used full length CAMTA3, suggesting the presence of domains that regulate DNA-binding. Future analysis of DNA binding should thus consider employing CAMTA3 fragments harbouring most of the phospho-sites but lacking these negatively-acting domains. Alternatively, one needs to investigate the requirement of Ca^2+^, CaM or other interacting proteins that may promote proper conformation of CAMTA3.

In this context, it is possible that CaM association to CAMTA3 alleviates DNA-binding. The relevance of CaM is further supported by the phenotype of the loss-of-function CAMTA3^K907E^ mutation (which is unable to bind CaM and does not repress expression of SA pathway genes) (Du et al., 2009) and the gain-of-function CAMTA3^A855V^ mutation (which is in a putative IQ motif) (Nie et al., 2012). Although no loss in CaM binding could be shown in CAMTA3^A855V^, this might be due to the presence of both IQ and CaMBD motifs and therefore quantitative differences may escape detection in the *in vitro* pull-down binding assays used. Alternatively, the A855V mutation affects regulation by CaM allosterically rather than changing binding affinity. Recent work in the Thomashow lab (Kim et al., 2017) showed that CAMTA3-mediated repression of defence genes involves an N-terminal repression module (NRM) that can act independently of CaM binding to the IQ or CaM-binding domains. While this challenges the prevalent concept of regulatory roles by CaM of CAMTA3, a possible explanation is that the partial CAMTA3 fragment (NRM) used by them contains the CAMTA3 DNA-binding domain, which is functional as a repressor when directly bound to target promoters. Their subsequent analysis using mutated full-length protein also showed that the suppressor function of the A855V mutation was dominant over the K907E mutation (i.e. CaM association at A855 can override loss of CaM association to the CaM-binding domain around K907). This indicates that there is differential regulation of NRM’s function through the IQ and CaM-binding domains. One may propose that the repressor function of the NRM domain alone is due to the uncoupling from the CaM regulation, thereby allowing it to bind DNA. Furthermore, Kim *et al*. (2017) also reported phosphorylation at S454 and/or S964 to partially contribute to suppressor function of full-length CAMTA3. S454 is one of the major MAPK-targeted sites identified in our report here. This strengthens the idea of regulation of CAMTA3 repressor function through phosphorylation. Unfortunately, we did not detect non-MAPK phospho-sites here, possibly reflecting different dynamics or sequential phosphorylation of CAMTA3, which were not addressed in the current study. Phosphorylation states at individual sites of the Elk-1 transcription factor have been shown to have opposing effects on transcriptional activation regulation after sequential modification by MAPKs (Mylona et al., 2016). Such multistep phosphorylation can act as tunable signalling circuits and therefore deserves attention in the future. However, the multiple phosphosites, possibly by sequential phosphorylation through MAPK or additionally a second kinase, will complicate the dissection of possible differential effects.

How CAMTA3 functions as a transcriptional repressor (Du et al., 2009; Nie et al., 2012) or an activator (Doherty et al., 2009) in different context is still unknown. A recent study questions the repressor function of CAMTA3 in defence-related genes (e.g. *EDS1*/*NDR1*). The authors proposed that the constitutive immune activation in *camta3* mutants is caused by activation of two NLRs, DSC1 and DSC2, and not caused by loss of CAMTA3 as a transcriptional suppressor (Lolle et al., 2017). While it is plausible that CAMTA3 is indeed a guardee of DSC1 and DSC2 and trigger immunity, the transcriptional repression activity of CAMTA3 involved in plant immunity in several other studies cannot be ignored. Foremost, the effects on immunity regulation is not restricted to the *camta3* knock-out mutants. CAMTA3 transgenic overexpression lines display compromised SAR and basal defence resistance (enhanced susceptibility to *Pst* DC3000) (Jing et al., 2011), but also reduced basal expression level of defence-related genes (*EDS1* and *NHL10*) and flg22-induced expression level of these genes (Fig 1). Furthermore, phenotypes of the CAMTA3^A855V^ gain-of-function mutant (Jing et al., 2011; Nie et al., 2012) or the loss-of-function CAMTA3^K907E^ (Du et al., 2009) cannot be explained by guarding through DSC1/DSC2 since CAMTA3 proteins are present. A possible scenario would be that DSC1/DSC2 is not guarding the absence of CAMTA3 but rather may be monitoring a molecular process or a component involved in the CAMTA3-CaM interaction via the IQ or CaM-binding motifs.

## Conclusions

In conclusion, the isolation of *CAMTA3* mutants in numerous independent genetic screens for pathogen/stress responses coupled with its post-translational control through (direct/indirect) phosphorylation by two major stress-responsive kinase pathways, namely MAPKs and CDPKs, highlights it as a node in abiotic and biotic stress signalling network. The convergence between environmental (cold) and pathogen (or other biotic stresses) signalling on CAMTA3 suggests it may control the interplay and trade-offs between growth and stress response. There are still many gaps in the understanding of how CAMTA3 navigates these different roles but multiple phosphorylation and nucleo-cytoplasmic trafficking may be involved.

## Author contributions

XJ, DS, JL conceived and designed the experiments. XJ performed the biochemical experiments and WH analyzed the phosphopeptide identification data. XJ & JL wrote the manuscript with input from all co-authors.

## Acknowledgements

We thank Prof. Hillel Fromm and Prof. Tina Romeis for discussions and suggestions within the scope of our GIF-funded collaborative project. Dr Anja Liese and Dr Marie Boudsouq kindly provided various CPK constructs. The expression constructs for expression of CA-MAPKs were a kind gift from Dr. Jean Colcombert. We also thank Prof Yuelin Zhang for seeds of the *camta3-3D* mutant harbouring the A855V mutation. Samuel Grimm provided the positive controls for the split-LUC assay (cLUC-WRKY34 and MVQ1-nLUC). We further thank Nicole Bauer and members of the Proteome Analytics group (Petra Majovsky, Carsten Proksch and Domenika Thieme) for excellent technical support, and members of our group, particularly Dr Lennart Eschen-Lippold, for discussions and technical advices.

## Materials and Methods

### Accession numbers of key genes mentioned in this work

CAMTA3 (At2g22300), SR1IP1 (At5g67385), MPK3 (At3g45640), MPK6 (At2g43790), CPK5 (AT4G35310).

### Plant growth conditions, protoplast assays and immunoblot analysis

*Arabidopsis thaliana* plants were grown on soil in climate chambers for 5-6 weeks (22° C; 8 h of light, 16 h of darkness; 140 µE). Protoplast isolation and transfection were performed as described (Yoo et al., 2007). Proteins were extracted by directly adding SDS-loading buffer to the pelleted protoplasts and processed for immunoblotting as described (Lee et al., 2004).

### General molecular cloning

Coding sequences of genes used in transient expression experiments were subcloned in either pDONR221 or pENTR-D-Topo vectors (Thermo Fisher Scientific). Gateway LR Clonase II Enzyme mix (Thermo Fisher Scientific) was used to generate constructs in destination vectors. Primers and description of transient expression constructs used in this study are listed in Table S1 and S4. Site-directed mutagenesis was performed using the primers listed in Table S2 as previously described (Palm-Forster et al., 2012; Eschen-Lippold et al., 2014).

### Real-time PCR analysis

Total RNA was extracted from plant tissues using TRIzol reagent (Roth). 2 µg of RNA was used for cDNA synthesis (Thermo Fisher Scientific). Diluted cDNA (1:10) was used for real-time PCR following the manufacturer’s protocol from 5×QPCR Mix EvaGreen® (ROX) (Bio & Sell). Primers and probes are listed in Table S3. The PCR was performed in MX3005P cyclers (Agilent), and the program consisted of an initial activation step (95°C, 15 min) followed by 40 cycles (15 s at 95°C and 60 s at 60°C).

### Protein stability assay

After transfection and overnight incubation for protein expression, 300 µl of the protoplasts were elicited with 100 nM flg22 for the indicated time (H_2_O was used as a mock control). Where indicated, 1 µM cycloheximide (CHX) was included to block translation. For the proteasome inhibitor, 50 µM of MG115 (Serva) was used to pretreat protoplasts for 30 min before flg22 and CHX elicitation. Cold treatment involves replacement with ice-cold media and immediate transfer to an ice-bath and cold room (4°C). To evaluate effect of specific kinases on protein stability, CAMTA3 was co-expressed with 35S-promoter-driven expression of *MKK5*^*DD*^, *CPK5-VK* or other indicated *CDPK*s. Protoplasts were pelleted and boiled in Laemmli loading buffer, separated on SDS-PAGE and processed for western blotting.

### Protein phosphorylation-dependent mobility shift assay in Phos-Tag^TM^ gels

Protoplasts (300 µl) transfected with pUGW14*-CAMTA3* (to express HA-tagged CAMTA3) were elicited with 100 nM flg22 and harvested at the indicated time points. To dephosphorylate the proteins, λ-phosphatase was used to treat extracted proteins according to the manufacturer’s protocol of Lambda Protein Phosphatase (New England Biolabs). Samples were boiled in Laemmli loading buffer and a Phos-Tag^TM^-based western blot was performed to analyze protein mobility shift. For separation of CAMTA3, Phos-TagTM SDS-PAGE gel was prepared to contain 6% acrylamide mix, 100 µM MnCl_2_ and 50 µM Phos-Tag^TM^ AAL-107 (Fujifilm Wako Chemicals) reagent. After separating protein samples on the Phos-Tag^TM^ gel, Mn^2+^ from the gel was eliminated to improve blotting transfer efficiency. The gel was washed 3 times 30 min with transfer buffer containing 10 mM EDTA and one time with 1 mM EDTA. Subsequently, proteins were transferred in a wet blot apparatus onto Nitrocellulose member (Macherey Nagel) for 135 min at 100 V.

### Quantitative Western blot

Odyssey® CLx multiplex imaging system (Li-COR) was used for quantitative western blot. After blotting, nitrocellulose membrane was stained with REVERT^TM^ Total Protein Stain (Li-COR), imaged in the Odyssey® imaging system (700 nm channel) and quantified with Image Studio^TM^ Software (Li-COR). Membrane was blocked with Odyssey® Blocking Buffer (Li-COR) for 1 hour and subsequently incubated with primary antibody diluted in Odyssey^TM^ Blocking Buffer (0.2% Tween 20) for 1 hour. After a brief rinse, membrane was incubated with secondary antibody IRDye® 800CW Goat anti-Mouse or anti-Rabbit IgG (H+L) (Li-COR) diluted in Odyssey^TM^ Blocking Buffer (0.2% Tween 20) for 1 hour. After wash steps, target proteins were imaged (800 nm channel) as described above. To compensate for loading differences, the target protein signals (800 nm) were normalized against the total protein quantification (700 nm) according to the Odyssey® CLx Application Protocol Manual (Li-COR). Typically, three (or more) biological replicate experiments were performed. The normalized values from individual experiment were further subjected to median-polishing prior to statistical analysis.

### Recombinant protein expression and purification

CAMTA3-WT and phospho-mutant variants were expressed as His_10_-tagged fusion proteins via pDEST-N110 in KRX (Promega) *E. coli* cells. Protein expression was induced with 0.1% Rhamnose at 20°C for 10 hours. MPK3, MPK4 and MPK6 were expressed as GST-tagged fusion proteins via pGEX4T-1 in BL21 cells, and expression induced with 1 mM IPTG at 16°C for overnight. Standard Ni-NTA or GSH-sepharose purification procedures were followed.

### Protein immunoprecipitation from protoplasts

To obtain CPK5 or constitutively-active MPKs for kinase assays, protoplasts were transfected with the corresponding constructs and proteins were immunoprecipitated after an overnight expression period. The following plasmids were used: pEXSG encoding CPK5-FL-strepII or CPK5m-FL-strepII (Dubiella et al., 2013); p2HAGW7 encoding HA-tagged MAPKs (Berriri et al., 2012; Genot et al., 2017). For CPK5, protoplasts were lysed in a modified extraction buffer (100 mM Tris pH 8.0; 5 mM EGTA; 5 mM EDTA; 100 mM NaCl; 20 mM DTT; 0.5 mM AEBSF; 2 µg/ml Aprotinin; 2 µg/ml Leupeptin; Protease-Inhibitor Mix P (Serva); 0.5% TritonX-100), and immunoprecipitated with MagStrep “Type3” XT beads (IBA Lifesciences). For MAPKs, MAPK extraction buffer (25 mM Tris pH 7.8; 75 mM NaCl; 15 mM EGTA; 15 mM β-Glycerophosphate; 15 mM 4-Nitrophenylphosphate; 10 mM MgCl_2_; 1 mM NaF; 0.5 mM Na_3_VO_4_; 1 mM DTT; 0.1% Tween20; 1 mM PMSF; 10 µg/ml Leupeptin; 10 µg/ml Aprotinin) was used and immunoprecipitated with HA antibody coupled to Protein G Sepharose® 4 Fast Flow (GE Healthcare).

### *In vitro* kinase assay

*In vitro* phosphorylation assay of CAMTA3 by MAPKs was performed in kinase substrate buffer 1 (20 mM HEPES pH 7.5; 15 mM MgCl_2_; 5 mM EGTA; 1 mM DTT; Protease-Inhibitor Mix HP; 2 µCi [gamma-^32^P]ATP) using recombinant His-CAMTA3 variants and GST-MPK3, 4 and 6. Samples were incubated for 30 min at 30°C, and the reactions were stopped by adding 5× loading buffer and 5 min boiling. Samples were subsequently separated on SDS-PAGE, stained with CBB and analyzed by autoradiography. Similarly, i*n vitro* phosphorylation assay of CAMTA3 by strepII-CPK5-FL or His-tagged CPK5-VK was performed as above except that EGTA was excluded and replaced with 0.5 µg Calmodulin/0.1 mM CaCl_2_. Reciprocally, to check for calcium dependency (negative control), CaCl_2_ was replaced with 2.4 mM EGTA.

### Phosphorylation site mapping

Phosphorylated CAMTA3 was obtained by *in vitro* phosphorylation with MPK3 or MPK6, or immunoprecipitated from leaves of CAMTA3 overexpressing plants that were infiltrated with flg22 (1 µM, 1 h). Amino acid residue-specific phosphorylation of CAMTA3 was mapped by liquid chromatography on-line with high resolution accurate mass MS (HR/AM LC-MS). Proteins separated by SDS-PAGE were subjected to in-gel tryptic digestion or with a combination of Glu-C. The resulting peptides were separated using C18 reverse phase chemistry employing a pre-column (EASY column SC001, length 2 cm, ID 100 μm, particle size 5 μm) in line with an EASY column SC200 with a length of 10 cm, an inner diameter (ID) of 75 μm and a particle size of 3 μm on an EASY-nLC II (all from Thermo Fisher Scientific). Peptides were eluted into a Nanospray Flex ion source (Thermo Fisher Scientific) with a 60 or 120 min gradient increasing from 5% to 40% acetonitrile in ddH_2_O with a flow rate of 300 nl/min and electrosprayed into an Orbitrap Velos Pro mass spectrometer (Thermo Fisher Scientific). The source voltage was set to 1.9 kV, the S Lens RF level to 50%. The delta multipole offset was −7.00. The AGC target value was set to 1e06 and the maximum injection time (max IT) to 500 ms in the Orbitrap. The parameters were set to 1e04 and 100 ms in the LTQ with an isolation width of 2 Da for precursor isolation and MS/MS scanning. Peptides were analyzed by a targeted data acquisition (TDA) scan strategy with inclusion list to specifically select and isolate CAMTA3 phosphorylated peptides for MS/MS peptide sequencing. Multi stage activation (MSA) was applied to further fragment ion peaks resulting from neutral loss of the phosphate moiety by dissociation of the high energy phosphate bond to generate b- and y-fragment ion series rich in peptide sequence information.

MS/MS spectra were used to search the TAIR10 database (ftp://ftp.arabidopsis.org, 35394 sequences, 14486974 residues) with the Mascot software v.2.5 linked to Proteome Discoverer v.1.4. The enzyme specificity was set to trypsin and two missed cleavages were tolerated. Carbamidomethylation of cysteine was set as a fixed modification and oxidation of methionine and phosphorylation of serine and threonine as variable modifications. The precursor tolerance was set to 7 ppm and the product ion mass tolerance was set to 0.8 Da. A decoy database search was performed to determine the peptide spectral match (PSM) and peptide identification false discovery rates (FDR). Phosphorylated peptides with a score surpassing the false discovery rate threshold of 0.01 (q-value<0.01) were considered positive identifications. The phosphoRS module was used to specifically map phosphorylation to amino acid residues within the primary structure of phosphopeptides.

### Microscopy

For localization studies, pUBC-based plasmids (Grefen et al., 2010) expressing *CAMTA3-YFP* or *SR1IP1-CFP* fusions were transfected into protoplasts. Where indicated, pEXSG-*ERF104-CFP* (Bethke et al., 2009) was included as a nuclear marker for co-localization studies or pRT100myc*-MKK5*^*DD*^ (or - *MKK5*^*KR*^) (Lee et al., 2004) were co-transfected for in vivo activation of MPK3/6. For BiFC assay, protoplasts were co-transfected with constructs of pE-SPYNE/pUC-SPYNE or pE-SPYCE/pUC-SPYCE (Walter et al., 2004) coding indicated pairs to be analyzed (see Table S4 for details). After an overnight incubation, protoplasts were directly analyzed by confocal laser scanning microscopy with an LSM780 (Zeiss). Fluorescence image settings are: YFP (Ex: 514 nm, Em: 500-570 nm) or CFP (Ex: 458 nm, Em: 480-520 nm).

### Statistical analyses

Statistical significance was analyzed with Prism 5 (GraphPad) software.

## Supplemental Information

Figure S1 MS2 spectra of peptides with phosphorylated site found in CAMTA3.

Figure S2 Yeast- or plant-based interaction assays between CAMTA3 and SR1IP1.

Figure S3 Flg22 treatment induces subcellular re-localization of CAMTA3.

Figure S4 EMSA for *in vitro* binding of full-length CAMTA3 to a “CGCG-containing” promoter fragment of EDS1.

Figure S5 CAMTA3 is not phosphorylated by CPK5 *in vitro*.

Figure S6 Compilation of western blots validating the expression of intact fusion proteins for the BiFC or co-localization experiments.

Table S1: Primers for cloning into entry vector pENTR/D

Table S2: Primers used for site-directed mutagenesis of CAMTA3 in entry vectors

Table S3: Primers and probes for RT-qPCR

Table S4: List of constructs used in this study

## References

Asai T, Tena G, Plotnikova J, Willmann MR, Chiu WL, Gómez-Gómez L, Boller T, Ausubel FM, Sheen J (2002) MAP kinase signalling cascade in Arabidopsis innate immunity. Nature 415: 977–983

Benn G, Wang CQ, Hicks DR, Stein J, Guthrie C, Dehesh K (2014) A key general stress response motif is regulated non-uniformly by CAMTA transcription factors. Plant J 80: 82–92

Berriri S, Garcia AV, Frei dit Frey N, Rozhon W, Pateyron S, Leonhardt N, Montillet JL, Leung J, Hirt H, Colcombet J (2012) Constitutively active mitogen-activated protein kinase versions reveal functions of Arabidopsis MPK4 in pathogen defense signaling. Plant Cell 24: 4281–4293

Bethke G, Pecher P, Eschen-Lippold L, Tsuda K, Katagiri F, Glazebrook J, Scheel D, Lee J (2012) Activation of the Arabidopsis thaliana mitogen-activated protein kinase MPK11 by the flagellin-derived elicitor peptide, flg22. Mol Plant Microbe Interact 25: 471–480

Bethke G, Unthan T, Uhrig JF, Pöschl Y, Gust AA, Scheel D, Lee J (2009) Flg22 regulates the release of an ethylene response factor substrate from MAP kinase 6 in *Arabidopsis thaliana* via ethylene signaling. Proc Natl Acad Sci U S A 106: 8067–8072

Boudsocq M, Sheen J (2013) CDPKs in immune and stress signaling. Trends in plant science 18: 30–40

Boudsocq M, Willmann MR, McCormack M, Lee H, Shan L, He P, Bush J, Cheng SH, Sheen J (2010) Differential innate immune signalling via Ca^2+^ sensor protein kinases. Nature 464: 418–422

Cheval C, Aldon D, Galaud J-P, Ranty B (2013) Calcium/calmodulin-mediated regulation of plant immunity. Biochimica et Biophysica Acta (BBA) - Molecular Cell Research 1833: 1766–1771

Chinchilla D, Zipfel C, Robatzek S, Kemmerling B, Nürnberger T, Jones JD, Felix G, Boller T (2007) A flagellin-induced complex of the receptor FLS2 and BAK1 initiates plant defence. Nature 448: 497–500

DeFalco Thomas A, Bender Kyle W, Snedden Wayne A (2010) Breaking the code: Ca^2+^ sensors in plant signalling. Biochemical Journal 425: 27–40

Doherty CJ, Van Buskirk HA, Myers SJ, Thomashow MF (2009) Roles for Arabidopsis CAMTA transcription factors in cold-regulated gene expression and freezing tolerance. Plant Cell 21: 972–984

Du L, Ali GS, Simons KA, Hou J, Yang T, Reddy AS, Poovaiah BW (2009) Ca^2+^/calmodulin regulates salicylic-acid-mediated plant immunity. Nature 457: 1154–1158

Du L, Ali GS, Simons KA, Hou J, Yang T, Reddy ASN, Poovaiah BW (2009) Ca2+/calmodulin regulates salicylic-acid-mediated plant immunity. Nature 457: 1154–1158

Dubiella U, Seybold H, Durian G, Komander E, Lassig R, Witte CP, Schulze WX, Romeis T (2013) Calcium-dependent protein kinase/NADPH oxidase activation circuit is required for rapid defense signal propagation. Proc Natl Acad Sci U S A 110: 8744–8749

Eschen-Lippold L, Bauer N, Lohr J, Palm-Forster MA, Lee J (2014) Rapid mutagenesis-based analysis of phosphorylation sites in mitogen-activated protein kinase substrates. Methods Mol Biol 1171: 183–192

Eschen-Lippold L, Bethke G, Palm-Forster MA, Pecher P, Bauer N, Glazebrook J, Scheel D, Lee J (2012) MPK11-a fourth elicitor-responsive mitogen-activated protein kinase in *Arabidopsis thaliana*. Plant Signal Behav 7: 1203–1205

Finkler A, Ashery-Padan R, Fromm H (2007) CAMTAs: Calmodulin-binding transcription activators from plants to human. FEBS Letters 581: 3893–3898

Galon Y, Nave R, Boyce JM, Nachmias D, Knight MR, Fromm H (2008) Calmodulin-binding transcription activator (CAMTA) 3 mediates biotic defense responses in Arabidopsis. FEBS Lett 582: 943–948

Gao X, Chen X, Lin W, Chen S, Lu D, Niu Y, Li L, Cheng C, McCormack M, Sheen J, Shan L, He P (2013) Bifurcation of Arabidopsis NLR immune signaling via Ca^2+^-dependent protein kinases. PLoS Pathog 9: e1003127

Genot B, Lang J, Berriri S, Garmier M, Gilard F, Pateyron S, Haustraete K, Van Der Straeten D, Hirt H, Colcombet J (2017) Constitutively Active Arabidopsis MAP Kinase 3 Triggers Defense Responses Involving Salicylic Acid and SUMM2 Resistance Protein. Plant Physiol 174: 1238–1249

Gómez-Gómez L, Boller T (2000) FLS2: an LRR receptor-like kinase involved in the perception of the bacterial elicitor flagellin in Arabidopsis. Mol Cell 5: 1003–1011

Grefen C, Donald N, Hashimoto K, Kudla J, Schumacher K, Blatt MR (2010) A ubiquitin-10 promoter-based vector set for fluorescent protein tagging facilitates temporal stability and native protein distribution in transient and stable expression studies. Plant J 64: 355–365

Han L, Li GJ, Yang KY, Mao G, Wang R, Liu Y, Zhang S (2010) Mitogen-activated protein kinase 3 and 6 regulate *Botrytis cinerea*-induced ethylene production in Arabidopsis. Plant J 64: 114–127

Heazlewood JL, Durek P, Hummel J, Selbig J, Weckwerth W, Walther D, Schulze WX (2008) PhosPhAt: a database of phosphorylation sites in Arabidopsis thaliana and a plant-specific phosphorylation site predictor. Nucleic Acids Research 36: D1015–D1021

Hoehenwarter W, Thomas M, Nukarinen E, Egelhofer V, Röhrig H, Weckwerth W, Conrath U, Beckers GJ (2012) Identification of novel *in vivo* MAP kinase substrates in *Arabidopsis thaliana* through use of tandem metal oxide affinity chromatography. Mol Cell Proteomics 12: 369–380

Hruz T, Laule O, Szabo G, Wessendorp F, Bleuler S, Oertle L, Widmayer P, Gruissem W, Zimmermann P (2008) Genevestigator v3: a reference expression database for the meta-analysis of transcriptomes. Adv Bioinformatics 2008: 420747

Jacob F, Kracher B, Mine A, Seyfferth C, Blanvillain-Baufumé S, Parker JE, Tsuda K, Schulze-Lefert P, Maekawa T (2018) A dominant-interfering camta3 mutation compromises primary transcriptional outputs mediated by both cell surface and intracellular immune receptors in Arabidopsis thaliana. New Phytologist 217: 1667–1680

Jing B, Xu S, Xu M, Li Y, Li S, Ding J, Zhang Y (2011) Brush and spray: a high-throughput systemic acquired resistance assay suitable for large-scale genetic screening. Plant Physiol 157: 973–980

Kadota Y, Sklenar J, Derbyshire P, Stransfeld L, Asai S, Ntoukakis V, Jones JD, Shirasu K, Menke F, Jones A, Zipfel C (2014) Direct regulation of the NADPH oxidase RBOHD by the PRR-associated kinase BIK1 during plant immunity. Mol Cell 54: 43–55

Kamiyoshihara Y, Iwata M, Fukaya T, Tatsuki M, Mori H (2010) Turnover of LeACS2, a wound-inducible 1-aminocyclopropane-1-carboxylic acid synthase in tomato, is regulated by phosphorylation/dephosphorylation. Plant J 64: 140–150

Kim YS, An C, Park S, Gilmour SJ, Wang L, Renna L, Brandizzi F, Grumet R, Thomashow M (2017) CAMTA-Mediated Regulation of Salicylic Acid Immunity Pathway Genes in Arabidopsis Exposed to Low Temperature and Pathogen Infection. Plant Cell

Kinoshita E, Kinoshita-Kikuta E, Koike T (2009) Separation and detection of large phosphoproteins using Phos-tag SDS-PAGE. Nat Protoc 4: 1513–1521

Laluk K, Prasad KV, Savchenko T, Celesnik H, Dehesh K, Levy M, Mitchell-Olds T, Reddy AS (2012) The calmodulin-binding transcription factor SIGNAL RESPONSIVE1 is a novel regulator of glucosinolate metabolism and herbivory tolerance in Arabidopsis. Plant Cell Physiol 53: 2008–2015

Lassowskat I, Bottcher C, Eschen-Lippold L, Scheel D, Lee J (2014) Sustained mitogen-activated protein kinase activation reprograms defense metabolism and phosphoprotein profile in Arabidopsis thaliana. Frontiers in Plant Science 5

Lee J, Eschen-Lippold L, Lassowskat I, Bottcher C, Scheel D (2015) Cellular reprogramming through mitogen-activated protein kinases. Frontiers in Plant Science 6

Lee J, Rudd JJ, Macioszek VK, Scheel D (2004) Dynamic changes in the localization of MAPK cascade components controlling pathogenesis-related (PR) gene expression during innate immunity in parsley. Journal of Biological Chemistry 279: 22440–22448

Li L, Li M, Yu L, Zhou Z, Liang X, Liu Z, Cai G, Gao L, Zhang X, Wang Y, Chen S, Zhou J-M (2014) The FLS2-associated kinase BIK1 directly phosphorylates the NADPH Oxidase RbohD to control plant immunity. Cell Host & Microbe 15: 329–338

Liu YD, Zhang SQ (2004) Phosphorylation of 1-aminocyclopropane-1-carboxylic acid synthase by MPK6, a stress-responsive mitogen-activated protein kinase, induces ethylene biosynthesis in Arabidopsis. Plant Cell 16: 3386–3399

Lolle S, Greeff C, Petersen K, Roux M, Jensen MK, Bressendorff S, Rodriguez E, Somark K, Mundy J, Petersen M (2017) Matching NLR Immune Receptors to Autoimmunity in camta3 Mutants Using Antimorphic NLR Alleles. Cell Host Microbe 21: 518–529 e514

Ludwig AA, Romeis T, Jones JD (2004) CDPK-mediated signalling pathways: specificity and cross-talk. J Exp Bot 55: 181–188

Lyzenga WJ, Stone SL (2012) Regulation of ethylene biosynthesis through protein degradation. Plant signaling & behavior 7: 1438–1442

Mylona A, Theillet FX, Foster C, Cheng TM, Miralles F, Bates PA, Selenko P, Treisman R (2016) Opposing effects of Elk-1 multisite phosphorylation shape its response to ERK activation. Science 354: 233–237

Nie H, Zhao C, Wu G, Wu Y, Chen Y, Tang D (2012) SR1, a calmodulin-binding transcription factor, modulates plant defense and ethylene-induced senescence by directly regulating NDR1 and EIN3. Plant Physiol 158: 1847–1859

Nitta Y, Ding P, Zhang Y (2014) Identification of additional MAP kinases activated upon PAMP treatment. Plant Signal Behav. 9: e976155

Palm-Forster MA, Eschen-Lippold L, Lee J (2012) A mutagenesis-based screen to rapidly identify phosphorylation sites in mitogen-activated protein kinase substrates. Anal Biochem 427: 127–129

Ranf S, Eschen-Lippold L, Pecher P, Lee J, Scheel D (2011) Interplay between calcium signalling and early signalling elements during defence responses to microbe- or damage-associated molecular patterns. Plant Journal 68: 100–113

Ranty B, Aldon D, Galaud J-P (2006) Plant Calmodulins and Calmodulin-Related Proteins: Multifaceted Relays to Decode Calcium Signals. Plant Signaling & Behavior 1: 96–104

Rayapuram N, Bonhomme L, Bigeard J, Haddadou K, Przybylski C, Hirt H, Pflieger D (2014) Identification of novel PAMP-triggered phosphorylation and dephosphorylation events in Arabidopsis thaliana by quantitative phosphoproteomic analysis. J Proteome Res 13: 2137–2151

Roux M, Schwessinger B, Albrecht C, Chinchilla D, Jones A, Holton N, Malinovsky FG, Tor M, de Vries S, Zipfel C (2011) The Arabidopsis leucine-rich repeat receptor-like kinases BAK1/SERK3 and BKK1/SERK4 are required for innate immunity to hemibiotrophic and biotrophic pathogens. Plant Cell 23: 2440–2455

Sebastià CH, Hardin SC, Clouse SD, Kieber JJ, Huber SC (2004) Identification of a new motif for CDPK phosphorylation in vitro that suggests ACC synthase may be a CDPK substrate. Archives of Biochemistry and Biophysics 428: 81–91

Seybold H, Trempel F, Ranf S, Scheel D, Romeis T, Lee J (2014) Ca2+ signalling in plant immune response: from pattern recognition receptors to Ca2+ decoding mechanisms. New Phytologist 204: 782–790

Thomma BP, Nürnberger T, Joosten MH (2011) Of PAMPs and effectors: the blurred PTI-ETI dichotomy. Plant Cell 23: 4–15

Walter M, Chaban C, Schütze K, Batistic O, Weckermann K, Näke C, Blazevic D, Grefen C, Schumacher K, Oecking C, Harter K, Kudla J (2004) Visualization of protein interactions in living plant cells using bimolecular fluorescence complementation. The Plant Journal 40: 428–438

Yang T, Poovaiah BW (2002) A Calmodulin-binding/CGCG Box DNA-binding Protein Family Involved in Multiple Signaling Pathways in Plants. Journal of Biological Chemistry 277: 45049–45058

Yoo SD, Cho YH, Sheen J (2007) Arabidopsis mesophyll protoplasts: a versatile cell system for transient gene expression analysis. Nat Protoc 2: 1565–1572

Zhang L, Du L, Shen C, Yang Y, Poovaiah BW (2014) Regulation of plant immunity through ubiquitin-mediated modulation of Ca(2+) -calmodulin-AtSR1/CAMTA3 signaling. Plant J 78: 269–281

Zhang T, Chen S, Harmon AC (2016) Protein-protein interactions in plant mitogen-activated protein kinase cascades. J Exp Bot 67: 607–618

Zipfel C, Robatzek S, Navarro L, Oakeley EJ, Jones JD, Felix G, Boller T (2004) Bacterial disease resistance in Arabidopsis through flagellin perception. Nature 428: 764–767

